# In Search for Biomarkers Reflecting Neural Implant-Induced Tissue Response Dynamics

**DOI:** 10.64898/2026.03.19.712876

**Authors:** Ali Sharbatian, Kevin Joseph, Ulrich G. Hofmann, Volker A. Coenen, Thomas Stieglitz, Danesh Ashouri

## Abstract

Extracellular matrix (ECM) remodeling is a fundamental determinant of neural tissue repair and implant integration, yet its conserved regulatory architecture remains undefined. While transcriptomic alterations following neural injury and implantation have been described, the ECM-centered programs that unify traumatic injury and neural implant responses remain unclear. Here, integrative systems-level transcriptomic analysis identifies a dominant and conserved ECM regulatory axis linking traumatic brain injury (BI), spinal cord injury (SCI), and neural implant–induced injury.

By integrating transcriptomic datasets from brain and spinal cord injury models using weighted gene co-expression network analysis (WGCNA), six conserved ECM-associated gene modules are identified, with hyaluronan (HA)-centered networks emerging as the dominant and conserved regulatory axis across both injury types. Modules enriched for low-molecular-weight HA (LMW-HA) are linked to Toll-like receptor signaling and pro-inflammatory cytokine expression, whereas high-molecular-weight HA (HMW-HA)–associated modules correlate with Cd44 signaling, tissue stabilization and repair.

Furthermore, independent validation in thin-film intracortical microelectrode datasets confirms robust activation of HA damage-associated molecular pattern (HA-DAMP) signaling following implantation, with 9/10 injury-derived modules preserved and 88% of transcripts exhibiting resolving temporal dynamics. These findings indicate that neural implants engage conserved trauma-associated ECM programs rather than a conventional foreign-body response, highlighting HA-related metabolisms. Given that HA fragments and HA-modifying enzymes are detectable in cerebrospinal fluid and peripheral circulation, HA-associated signatures may serve as minimally invasive biomarkers of neural injury and implant biocompatibility, enabling longitudinal monitoring and informing next-generation neural interface design.

## Introduction

Neural implants like intracortical micro electrodes (IMEs) enable direct communication with brain circuits and hold promise as potential tool for treatment and rehabilitation for a wide range of neurological conditions. However, their long-term performance remains constrained by complex tissue responses at the device–tissue interface, including variability in chronic unit yield and progressive signal loss (Kozai et al. 2015; Usoro et al. 2021). In the case of intracortical devices like depth electrodes, the initial insertion produces a localized microscopic traumatic brain injury (TBI), as shank devices penetrate the meninges, open the blood-brain barrier (BBB), generate shear forces, displace delicate parenchyma, and rupture microvessels (Otte et al. 2022). The resulting neurovascular trauma disrupts homeostasis and evokes innate immune activation that can extend beyond the immediate insertion site (Wellman et al. 2019; Potter et al. 2012).

In this work, we hypothesize that neural implant insertion activates conserved injury-response programs common to brain and spinal cord trauma, and we investigate whether these programs are reflected in distinct, quantifiable gene expression signatures associated with extracellular matrix (ECM) remodeling. While inflammation and gliosis are well described in implant contexts, the ECM has remained an underexplored integrator of mechanical injury, immune signaling, and long-term tissue remodeling. The ECM is a dynamic network of proteins and polysaccharides that regulates the structural integrity and signaling necessary for tissue maintenance and repair (Dityatev et al. 2010; Fawcett et al. 2019). Mechanical disruption of the ECM, in tandem with the blood–brain barrier (BBB) breach, and influx of serum proteins such as fibrinogen, albumin, and thrombin into parenchyma, are potent cofactors for microglial and astroglial activation (Takata et al. 2021; Kozai et al. 2015; Bouadi&Tay 2021). Peripheral immune cells, including neutrophils and lymphocytes, subsequently infiltrate brain tissue and amplify the inflammatory response (Bennett et al. 2018; Passaro et al. 2021). The breakdown of BBB integrity, coupled with neurovascular trauma, alters brain homeostasis and initiates a sustained immune activity in surrounding tissue (Wellman et al. 2019).

Unlike peripheral foreign body responses dominated by fibroblast-driven collagen deposition, reactions in the central nervous system (CNS) are governed by astrocytes and microglia, with contributions from infiltrating macrophages following BBB compromise (Adams und Gallo 2018; Salatino et al. 2017; Hoeferlin et al. 2025). The ensuing astrogliosis and matrix remodeling affect electrode-tissue coupling and recording stability, with variations depending on the implant type, implantation duration, and local tissue state. The ability of a neural implant to initiate, modulate and withstand these processes ultimately determines its biocompatibility.

As defined by the International Union of Pure and Applied Chemistry (IUPAC), biocompatibility describes ”the ability to interact with living systems without causing adverse effects.“ (Vert et al. 2012). ISO standards and regulatory agency approvals consider an implant biocompatible if a minimal scar capsule (50–200 microns) forms at the implant site 30 days after implantation, with minimal or no further reaction (Ratner 2016). While broadly applied, such static criteria fail to capture dynamic molecular processes that determine chronic neural interface performance, particularly those involving ECM components. Identifying the key molecular factors that shape wound healing, immune activation, and tissue remodeling represents a first step toward better assessing how a neural interface interacts with host tissue. Such insight can inform more rational design strategies aimed at improving device integration and long-term functional stability.

Linking neurovascular damage to immune signaling and local tissue remodeling the ECM appears to take a central role (Gaudet and Popovich 2014). Matrix remodeling can be organized along six principal ECM axes, each capturing distinct aspects of the injury response: (1) hyaluronan turnover and sensing, reflecting molecular weight–dependent signaling relevant to CNS injury; (2) provisional matrix, encompassing acute injury–response glycoproteins such as EDA fibronectin and tenascin-C; (3) perineuronal-net chondroitin sulfate proteoglycans (CSPGs), isolating the lectican family specific to CNS barrier function; (4) basement membrane components, representing neurovascular scaffolding including laminins, collagen IV, and perlecan; (5) proteases and regulators, including matrix metalloproteinases (MMPs), A Disintegrin and Metalloproteinase with Thrombospondin Motifs (ADAMTS), and Tissue Inhibitors of Metalloproteinases (TIMPs); and (6) crosslinking and fibrosis programmes, capturing scar-forming processes (Bradbury und Burnside 2019; Cooper et al. 2018).

ECM components are known to engage in distinct temporal and functional dynamics following neural injury. As example, following an injury, provisional matrix proteins, such as fibronectin and tenascin-C, are rapidly deposited (Tang et al. 2003), while chondroitin sulfate proteoglycans (CSPGs) accumulate within the glial scar and influence axonal regeneration (Silver und Miller 2004). Basement membrane elements are disrupted during BBB breakdown. Such mechanical or oxidative stress can act as damage-associated molecular patterns (DAMPs), activating innate immune receptors, including Toll-like receptors (Tlr2, Tlr4), Cd44, and the Cd14 coreceptor (Bedell et al. 2018; Jiang et al. 2007; Stern et al. 2006).

While these receptor–ligand interactions are well established in general injury contexts, their coordinated modular co-expression across multiple CNS injury types and their specific temporal dynamics in neural implant tissue have not been systematically characterized. Also, it remains unclear which ECM-associated axis predominates the case of neural implant injury response, and whether mechanisms identified in classical injury models translate to implant contexts. This knowledge gap limits our understanding of how the immune system “categorizes” a neural implant, whether as a transient injury that can be resolved or as a persistent stressor that drives chronic inflammation. A systematic dissection of ECM-centered injury programs may therefore reveal molecular signatures predictive of both neural injury as well as implant-induced injury.

To tackle this, we integrate differential gene expression and co-expression analysis with the aim of assessing major ECM-related dynamics and immune regulatory mechanisms. By systematically mining pathways governing ECM remodeling following neural injury, we aim to identify those associated with the tissue repair and immune modulation.

By leveraging existing datasets and published transcriptomic studies, we address two core objectives: First, we sought to identify ECM-related gene modules in neural injury through systematic WGCNA analysis, ranking ECM axes to determine which components show the most consistent activation. Second, we aim to test whether the top-ranked ECM axis, identified as trauma-activated in BI/SCI discovery models, is similarly engaged in neural implant tissue following insertion injury.

The study goes beyond state-of-the-art research on implant-related brain injury and unlike previous work that has examined inflammation or implant-induced damage in isolation, we hypothesize a regulatory mechanism in which ECM remodeling and immune activation are dynamically coordinated. This integrated perspective offers to open new opportunities for advancing biocompatibility assessment and supports the adoption of emerging diagnostic approaches, such as liquid biopsy-based (Abdelhak et al. 2022) analyses of biomarkers in peripheral fluids like blood for monitoring neural implant performance.

## Results

To investigate molecular responses underlying neural injuries, we conducted an integrative gene expression and functional annotation workflow (Figure 1). The resulting DEGs were visualized using volcano plots, to clearly depict upregulated and downregulated genes at different post-injury time points. Hierarchical clustering heatmaps further delineated gene expression patterns, clearly distinguishing between acute and chronic injury phases and highlighting coherent gene regulatory programs triggered by injury. Functional annotation through Gene Ontology (GO) and KEGG pathway analyses provided deeper biological insights, revealing that prominent biological processes included inflammatory signaling cascades, immune regulation, and extracellular matrix (ECM) remodeling. Hub genes identified through network analyses and their corresponding functional enrichment pathways are summarized explicitly in accompanying tables and graphical enrichment plots (Figure 1), collectively elucidating key molecular mechanisms activated following neural injury.

**Figure 1:**
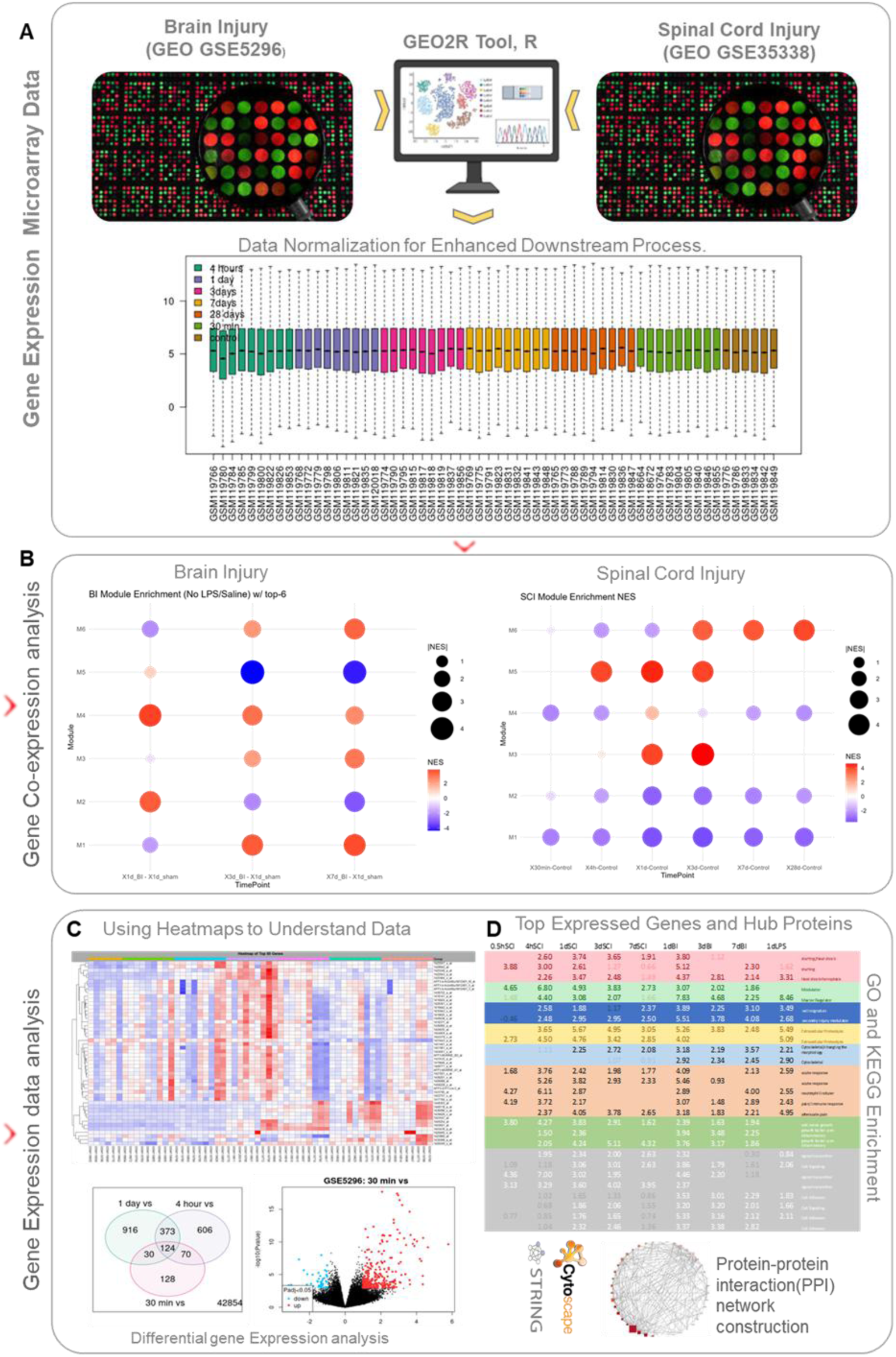
Workflow for gene expression analysis and functional annotation in neural injury models. (a) Gene expression microarray datasets were obtained from publicly available resources (GEO accession numbers GSE5296 for spinal cord injury and GSE35338 for brain injury). Data were processed and normalised using the GEO2R interface and R software. (b) WGC-NA identified enriched gene modules, visualised by normalised enrichment score (NES) bubble heatmaps, highlighting distinct temporal biological responses in brain and spinal cord injuries. (c) DEGs were identified (adjusted p-value < 0.05; log₂ fold change ≥ 1.0) and visualised via heatmaps, Venn diagrams, and volcano plots. (d) Hub proteins were identi-fied using a protein–protein interaction (PPI) network, and GO and KEGG analyses high-lighted critical processes.

### ECM Axis Ranking by Differentially Expressed Gene Counts

To characterize ECM remodeling after neural injury, differentially expressed genes (DEGs) were analyzed within our six ECM functional axes defined for this study (see Methods): (1) Hyaluronan turnover and sensing, (2) Provisional matrix (EDA-fibronectin and Tenascin-C), (3) Perineuronal net chondroitin sulfate proteoglycans (PNN-CSPGs), (4) Basement membrane, (5) Proteases and regulators, and (6) Crosslinking/fibrosis.

Across both BI and SCI datasets, all six axes were transcriptionally engaged after injury, but with distinct amplitudes, durations, and composition profiles. Axis-level enrichment (GSVA/ssGSEA) and DEG counts revealed three major temporal phases of ECM remodeling: (1) Early innate and provisional activation, dominated by hyaluronan and provisional-matrix axes; (2) Barrier remodeling, centered on basement-membrane and Protease activity; and (3) Matrix consolidation, characterized by crosslinking and fibrosis.

When ranked by the number of unique DEGs significant at one or more timepoints, the hyaluronan (HA) axis consistently showed the strongest and most reproducible activation (rank 1 in both BI and SCI; Table 2). In BI, the HA axes comprised five DEGs, followed by the Protease/Regulator and CSPG-PNN axes. In SCI, HA again ranked first with six DEGs, while the provisional, CSPG-PNN, and crosslinking/Fibrosis axes shared second rank.

**Table 1.**
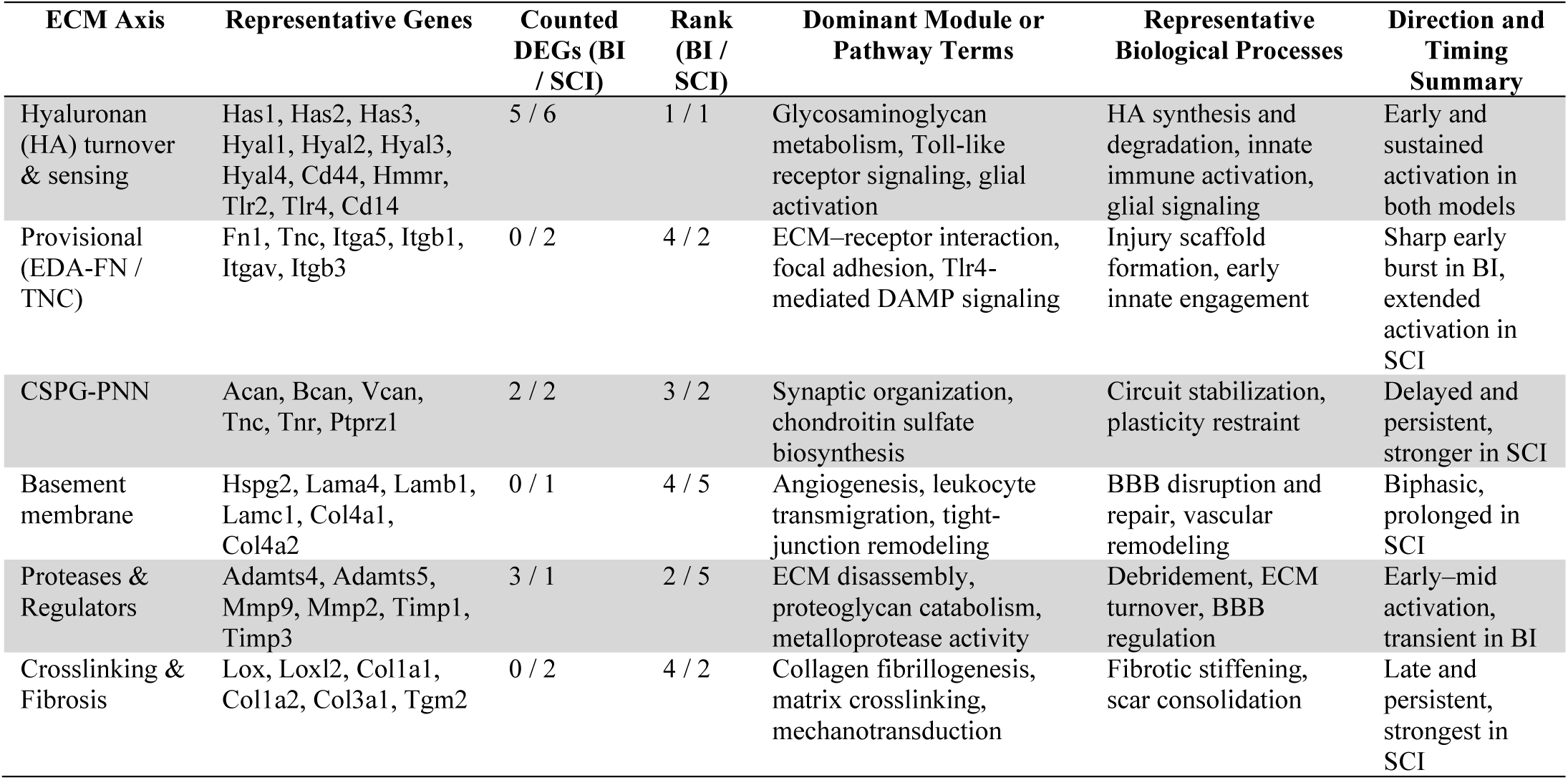
Axis-level DEG statistics and dominant functional themes across brain and spinal cord injury datasets.

**Table 2.**
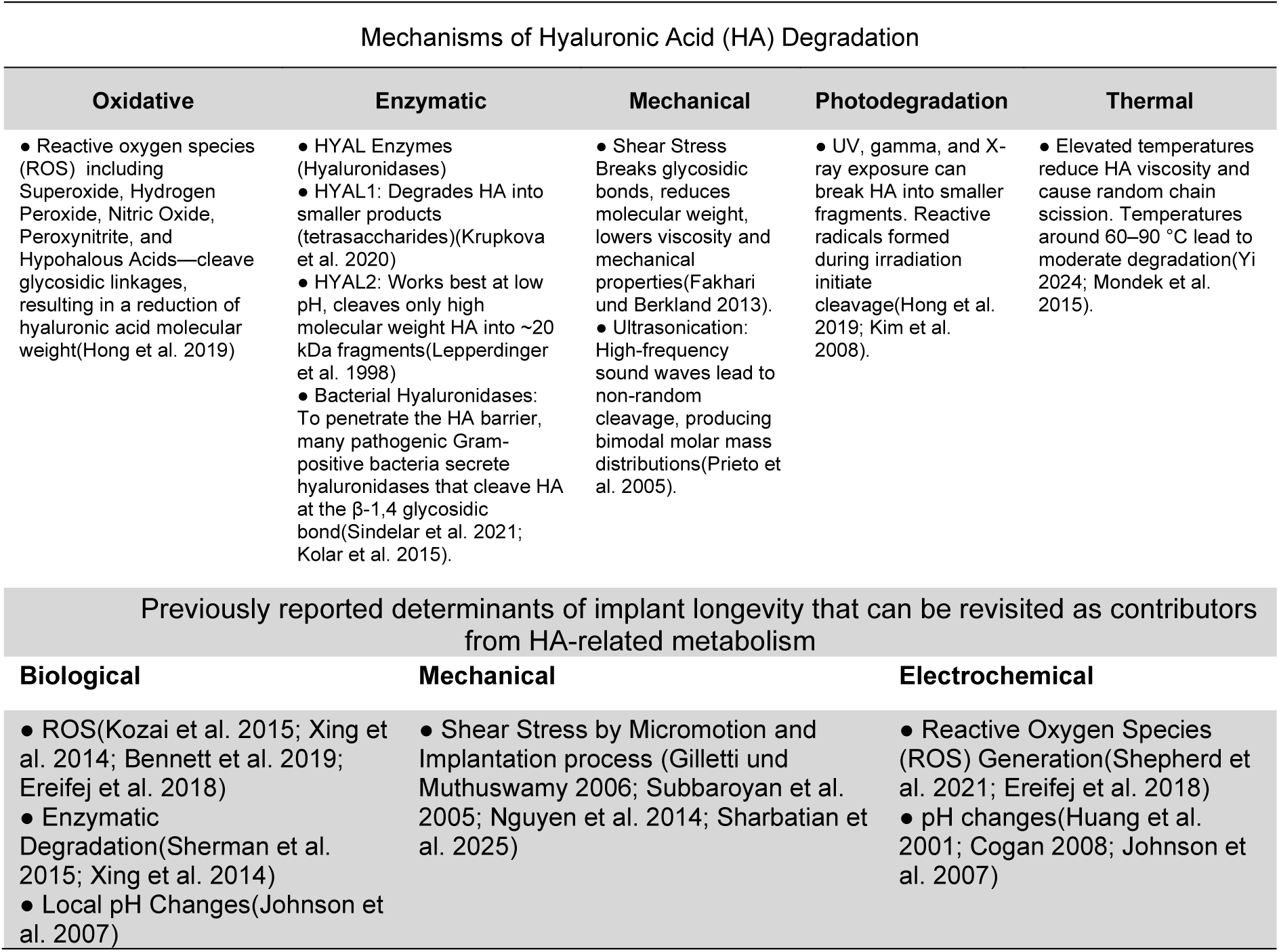
Mechanisms of Hyaluronic Acid (HA) degradation and implant-specific pathways. The upper panel summarizes the five principal routes of HA degradation (oxidative, enzymatic, mechanical, photodegradation, and thermal), together with key references. The lower panel maps these mechanisms to the three factor categories most relevant to intracortical microelectrode arrays: biological, mechanical, and electrochemical.

Together, these quantitative trends delineate a conserved HA-centered ECM remodeling program with injury-specific temporal dynamics. BI exhibited a compressed ECM response, defined by a rapid and coordinated HA–Provisional surge, limited protease activation, transient Basement-membrane remodeling, modest CSPG accumulation, and minimal fibrosis. In contrast, SCI showed a prolonged and amplified ECM response, characterized by sustained HA and Provisional activity, extended barrier remodeling, broader protease participation, prominent PNN-CSPG deposition, and persistent fibrotic consolidation.

The HA axis was enriched for glycosaminoglycan metabolism, Toll-like receptor signaling, and glial activation; the Provisional matrix axis was linked to ECM–receptor interaction, focal adhesion, and Tlr4-mediated DAMP signaling; the CSPG-PNN axis aligned with synaptic organization and chondroitin sulfate biosynthesis; the Basement-membrane axis reflected angiogenesis, leukocyte transmigration, and tight-junction remodeling; Protease/Regulator modules were associated with ECM disassembly and metalloprotease activity; and the Crosslinking/Fibrosis axis was enriched for collagen fibrillogenesis, Lox/Tgm2 crosslinking, and mechanotransduction pathways.

Although all axes were curated from published ECM and injury-biology literature, the results indicated the Hyaluronan axis emerging as the central axis of metabolic, structural, and immunologic remodeling. It encompasses enzymes responsible for synthesis and degradation (Has1–3, Hyal1–3, CEMIP, TMEM2, SPAM1) together with receptor and signaling components (Cd44, HMMR, Tlr2, Tlr4, Cd14), thereby linking matrix turnover to innate immune sensing and cell–ECM communication.

Mechanistically, HA turnover is suggested here to be tightly coupled with the activity of other ECM axes. HA fragments act as endogenous DAMPs, activating Tlr2/4 and inducing expression of EDA-fibronectin and Tenascin-C in the Provisional matrix axis. MMPs and ADAMTSs from the Protease/Regulator axis further enhance HA degradation and release matrix-bound cytokines, amplifying HA-Tlr signaling. Subsequent remodeling of laminin and collagen IV within the Basement-membrane axis and CSPG deposition at perineuronal nets reflects a transition from reactive to stabilizing ECM states. Prolonged HA–Cd44/RHAMM signaling then promotes LOX, LOXL2, and TGM2 expression, driving collagen crosslinking, matrix stiffening, and fibrotic encapsulation.

Together, these results establish the Hyaluronan axis as a highly conserved and quantitatively dominant component of ECM remodeling across distinct CNS injury models. It is consistently engaged during early innate activation, is associated with provisional scaffold formation, and is linked to the transition toward fibrotic stabilization. The consistent prominence of HA-related pathways across both BI and SCI supports a unified model in which hyaluronan metabolism serves as a key component associated with inflammation, tissue mechanics, and repair outcomes in the injured central nervous system.

Given that differential expression analysis revealed that the Hyaluronan axis contained the highest DEG count in both injury models (BI: 5 DEGs; SCI: 6 DEGs; ranked 1^st^ of six axes in both models; see Table 2 and S1 for full axis-by-axis DEG counts) subsequent mechanistic analyses focused on hyaluronan-associated pathways.

### ECM Axis Enrichment Across Injury-Associated WGCNA Modules (BI and SCI)

WGCNA identified 20 modules in BI and 18 modules in SCI. For detailed functional annotation, we focused on the six largest modules per dataset (M1-M6), which captured > 70% of assigned genes for pathway enrichment (detailed module-trait correlations - injury status - are reported in Supplementary Table S2). Each WGCNA-derived gene co-expression modules (Mi) was mapped onto study-defined ECM axes (Table 2) to systematically identify which ECM programs dominates the neural injury response.

The six largest modules (M1–M6) per injury models (see Supplementary Table S2) were identified and functionally annotated by integrating dominant GO and KEGG enrichments, leading-edge genes, normalized enrichment scores (NES), and temporal expression profiles. Each module was mapped to these axes to capture the full ECM landscape beyond HA and to resolve axis–axis couplings within and across time. This categorization framework condenses complex transcriptomic dynamics into biologically interpretable coherent units, resolves temporal engagement of each ECM program, and enables direct comparison between BI and SCI across injury phases Where relevant below, the dominant ECM axis (or axes) contributing to each module is indicated.

In BI, temporal NES profiles delineate distinct phases of ECM axis responses (Figure 2). An early inflammatory peak at day 1 (M2, Figure 2A–B), followed by sustained immune and translational activity from days 1-7 (M4) and, a late reparative/metabolic phase emerging at days 3-7 (M6). In parallel, neuronal and oxidative phosphorylation programs (M1) are suppressed from day 3 to day 7. Together, these patterns describe a BI trajectory that progresses from acute innate activation through prolonged glial/translational engagement to late-stage metabolic repair. Relative to SCI, BI shows a sharper early surge of the Hyaluronan and Provisional-matrix axes, a narrower window Basement-membrane remodeling window, and less persistent Crosslinking/fibrosis.

**Figure 2:**
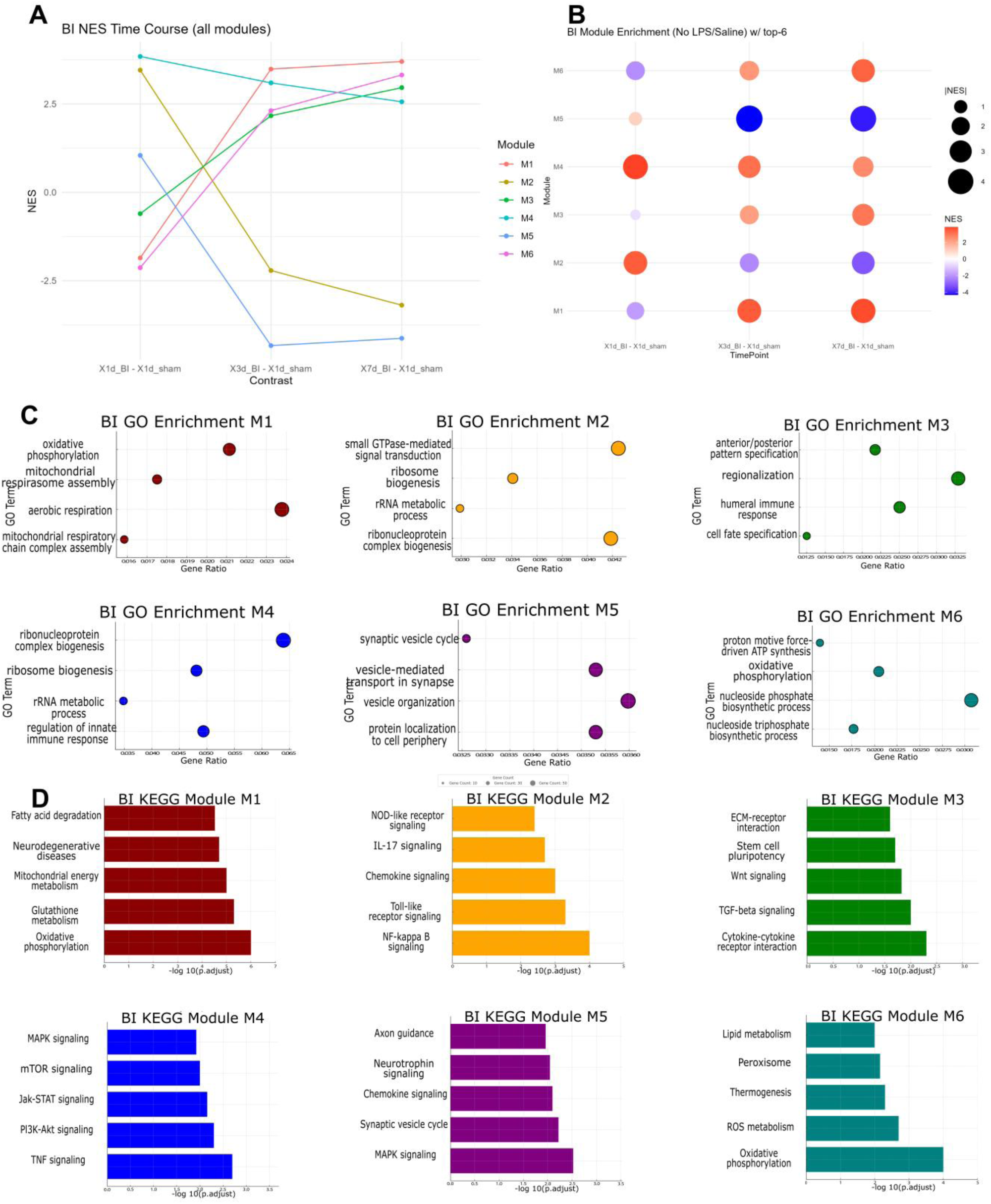
Functional and temporal characterization of ECM-associated WGCNA modules after brain injury. (A) Temporal profile of normalised enrichment scores (NES) for WGCNA modules (M1–M6) across post-injury time points (1 day, 3 days, and 7 days post-injury) relative to sham. Each line represents the dynamic activity of a module, highlighting distinct temporal patterns. (B) Bubble heatmap of NES values across time points for all modules. Bubble size reflects the significance of enrichment, and colour indicates the magnitude and direction of NES (red = upregulated, blue = downregulated), illustrating module-specific changes in activity over time. (C) GO biological process enrichment analysis for each module. Bubble plots display the top 4 significant GO terms per module. The x-axis shows the gene ratio (number of genes in the term/total module genes), the bubble size indicates gene count, and the colour corresponds to the module. (D) Top 5 enriched KEGG pathways for each module (M1–M6), shown as bar plots of –log₁₀(adjusted p-value). Bars are colour-coded by module, revealing distinct pathway-level functional associations. Panels also indicate, for each module, its dominant ECM axis mapping (hyaluronan, provisional, PNN-CSPG, basement membrane, proteases/regulators, crosslinking/fibrosis).

These succeeding phases map well onto HA biology: early low-molecular-weight HA (LMW-HA)–driven Tlr/NF-κB activation gives way to later high-molecular-weight HA (HMW-HA)–associated tissue stabilization and repair. When viewed through the ECM-axis lens, the same phases align with early provisional-matrix engagement (EDA-FN/TNC/integrins), intermediate basement-membrane remodeling around the BBB/NVU, timed protease activity (MMP/ADAMTS with TIMP regulation), and a modest late crosslinking/fibrosis signal.

Normalized Enrichment Scores (NES) were calculated across defined post-injury time points to quantify temporal dynamics. BI model samples were assessed at 1, 3, and 7 days post-injury, whereas SCI samples spanned 30 minutes, 4 hours, 1 day, 3 days, 7 days, and 28 days. A positive NES indicates that the genes in a given module are coordinately upregulated relative to sham or control samples, whereas a negative NES indicates coordinated downregulation. Throughout this section the terms upregulation and downregulation refer to these positive and negative NES shifts, respectively.

Analysis of NES temporal dynamics revealed distinct module expression trajectories, categorized into four primary response types:

- Transient activation – Module M2 (BI) peaks sharply at 1 d, falls by 3 d, and stabilises near baseline at day 7, consistent with a short-lived acute inflammatory burst (dominant ECM axes: hyaluronan; provisional matrix; early proteases/regulators).
- Persistent activation – Module M4 (BI) rises by day 1 and remains elevated at days 3 and 7, reflecting prolonged immune / protein-synthesis activity (dominant ECM axes: hyaluronan; provisional matrix; basement membrane repair; proteases/regulators).
- Delayed activation – Module M6 (BI) is quiescent at day 1, but climbs markedly at day 3 and is highest at day 7, matching late-stage reparative metabolism (dominant ECM axes: basement membrane maturation; PNN-CSPGs emergence; crosslinking/fibrosis low-to-moderate).
- Immediate and sustained suppression – Module M1 (SCI) drops steeply within 30 min and stays depressed through 28 d, indicating acute neuronal injury followed by long-term metabolic dysfunction (dominant ECM axes: PNN-CSPGs restraint co-occurs; basement membrane dysregulation early; crosslinking/fibrosis rises later).

Taken together, these NES profiles depict a finely graded molecular time-course for each injury model, underscoring the value of WGCNA for disentangling phase-specific gene-network activity and for pinpointing biomarkers appropriate to acute, sub-acute, and chronic stages of neural repair which offers valuable means to diagnostic procedures.

#### Module M1 (BI)

As the module associated with oxidative metabolism, M1 showed significant downregulation of neuronal oxidative phosphorylation and synaptic-transmission programs following BI. M1 expression declines sharply by day 3 (NES ≈ −3) and remained suppressed at day 7, consistent with profound neuronal suppression in the lesion area (Chen et al. 2013).

BI Module M1 indicated enriched in mitochondrial energy metabolism and reactive oxygen species (ROS) pathways. Top GO terms include mitochondrial ATP synthesis coupled to proton transport and oxidative phosphorylation, while KEGG pathways such as oxidative phosphorylation and glutathione metabolism are significantly over-represented. Hub genes identified included GRIA1 and CAMK2A, involved in synaptic plasticity, as well as myelin-related genes important for neural connectivity. M1 shows low Provisional and Protease activity internally, consistent with its neuronal identity; HA-turnover signals are instead concentrated in the acute inflammatory modules described below.

#### Module M2 (BI)

Module M2 (BI) exhibits a transient upregulation peakig at day 1 after BI, corresponding to the acute cytokine dynamics observed post-injury. Genes such as Il1b and Il6 surge within the first 24 hours and gradually decrease by days 3 and 7, mirroring kinetics of inflammatory cytokines like IL-1β and TNF-α.

Module M2 is strongly associated with innate immune and inflammatory responses, show enrichment in GO terms like positive regulation of innate immune response and inflammatory response. Key KEGG pathways identified include NF-κB, Toll-like receptor (Tlr), and chemokine signaling, underscoring the acute inflammatory nature of this module. In ECM-axis terms, M2 engages the Hyaluronan axis (HA fragment–driven TLR2/TLR4/ Cd14 signaling), Provisional matrix components (EDA-FN, TNC, integrin engagement), and early Protease/regulator activity.

#### Module M3 (BI)

Module 3 in BI demonstrates a moderate delayed upregulation, beginning after day 1 and persisting through day 7 post-injury. This upregulation pattern aligns with results in subacute phases of injury, characterized by microglia transitioning into phagocytic roles and reactive astrocytes initiating scar formation.

Module M3 pathways primarily involve cytokine signaling, cellular proliferation, and growth factor responses. The most significant KEGG pathway enrichment was cytokine–cytokine receptor interaction. Other enriched pathways included those regulating cell proliferation and stem cell pluripotency (e.g., TGF-β, Wnt, STAT3). Regarding ECM axes, M3 shows increased Protease/regulator activity (MMP/ADAMTS with TIMPs) and Basement-membrane repair signatures, while PNN-CSPG components (VCAN, ACAN, TNR) begin to accumulate. Elevated Has3 expression further indicates active hyaluronan synthesis.

#### Module M4 (BI)

Module 4 exhibits robust and sustained upregulation immediately after BI, reflecting the prolonged neuroinflammatory response characteristic of traumatic brain injury (TBI). The module was rapidly induced (NES significantly elevated at day 1) and remained persistently high at days 3 and 7 post-injury.

Gene expression patterns mirrored ongoing inflammatory processes, involving cytokines such as TNF-α, IL-6, CCL2, and astrocyte activation markers like GFAP. Gene Ontology (GO) analyses highlighted significant enrichment in ribosome biogenesis and RNA processing, indicating increased protein synthesis capability. Concurrently, inflammatory response processes (e.g., response to bacterial molecules, lipopolysaccharide-mediated responses, and inflammatory regulation) were notably active.

#### Module M5 (BI)

Module 5 demonstrated pronounced downregulation following BI, from day 1 to day 3 and remaining suppressed by day 7. The timeline matched histological observations of myelin disruption post-TBI, impaired oligodendrocyte survival and remyelination efforts. The module primarily encapsulated genes involved in intracellular vesicle trafficking, neurotransmitter secretion, exocytosis, and synaptic vesicle cycling, critical for neuronal communication and synaptic function. Diminished PNN-CSPG and Basement-membrane support within M5 parallels the observed synaptic and axonal suppression.

#### Module M6 (BI)

Module 6 was characterized by delayed and significant upregulation from day 3 to day 7, marking a transition toward repair and regeneration. This phase coincided with elevated expression of reparative growth factors such as TGF-β1, IGF-1, and angiogenesis signals like VEGF-A, as microglia and macrophages transitioned to a pro-repair phenotype.

Elevated Apoe expression further reflected lipid processing by macrophages/microglia from damaged myelin, supporting reparative and regenerative states. Module 6 prominently involved metabolic reprogramming, oxidative stress responses, and chromatin remodeling, as indicated by enriched KEGG pathways such as oxidative phosphorylation, thermogenesis, and reactive oxygen species (ROS) metabolism. Axis mapping: late basement-membrane repair, moderate PNN-CSPG assembly, and limited crosslinking/fibrosis co-occur with metabolic recovery.

Analyses driven from SCI displays immediate and sustained suppression of neuronal/oligodendrocyte modules (M1–M2; 30 min→ day 28), an early cytokine storm (M5; 4 h– day 1), prolonged antigen-processing/phagocytic activity (M3; 1–28 d), and late lipid/lysosomal metabolism (M6; 7–28 d) consistent with a more persistent fibrotic/ECM-remodeling response than BI (Figure 3). Across axes, SCI preserves early HA and Provisional signals longer, extends basement-membrane remodeling, amplifies protease/regulator breadth, builds stronger PNN-CSPG barriers, and culminates in durable crosslinking/fibrosis.

**Figure 3:**
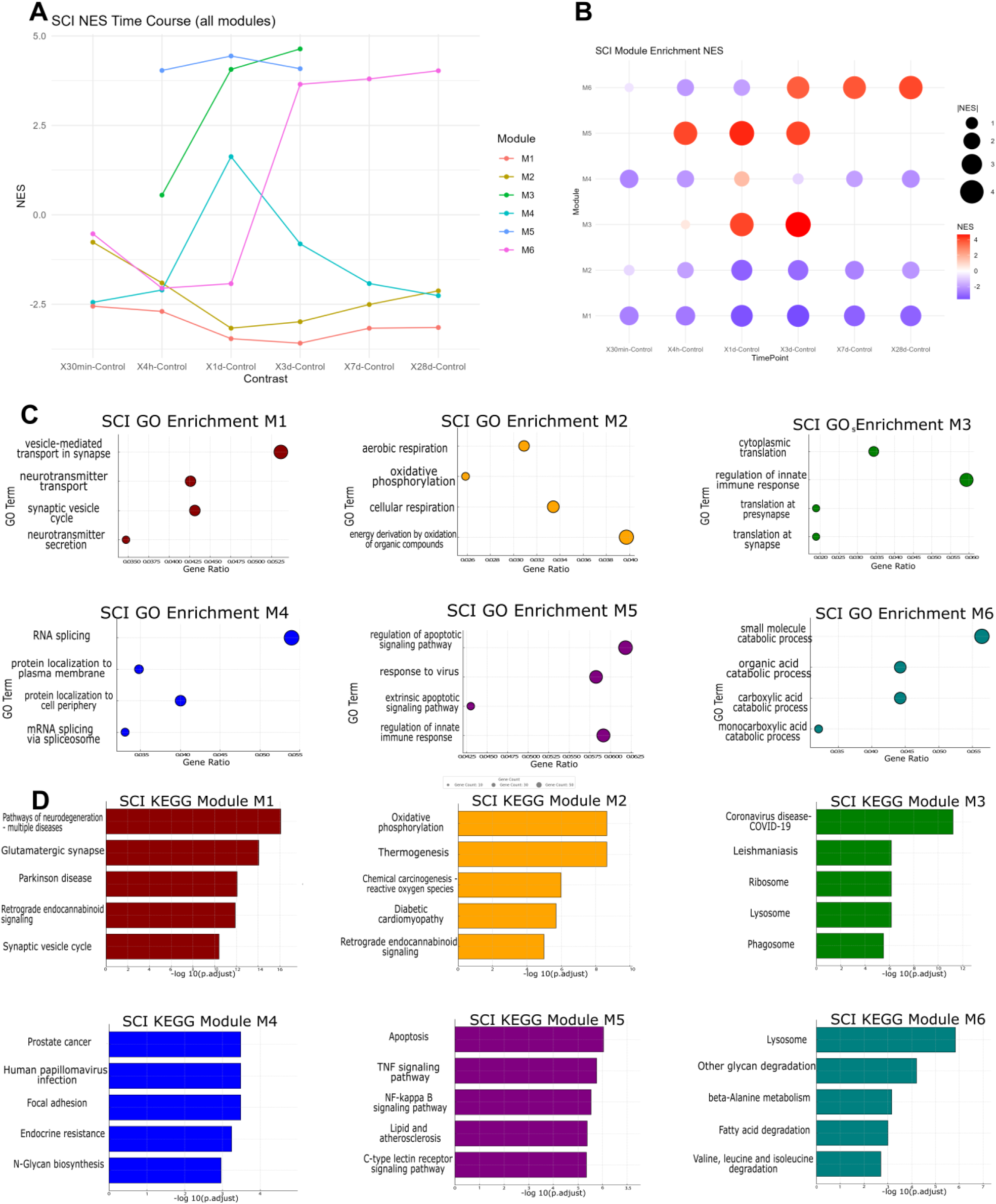
Functional and temporal characterization of SCI-associated WGCNA modules following spinal cord injury. (A) Temporal trajectory plots of normalized enrichment scores (NES) for WGCNA modules (M1–M6) across post-injury timepoints (30 minutes to 28 days post-SCI) compared to uninjured controls. Each line represents a module’s dynamic activity, capturing early and late transcriptional responses. (B) NES bubble heatmap summarizing temporal module activation patterns. Bubble color indicates NES direction and magnitude (red = activation, blue = suppression), while size reflects enrichment significance. This panel highlights temporally resolved functional shifts in gene regulation across SCI recovery phases. (C) GO biological process enrichment bubble plots for each module. The x-axis shows gene ratio (enriched genes per total module size), bubble size reflects gene count, and color denotes the module. Top 4 GO terms are shown per module, highlighting distinct biological processes such as mitochondrial function, immune response, and lipid metabolism. (D) Top 5 enriched KEGG pathways for each SCI-associated module (M1–M6), visualized as –log₁₀(adjusted p-value) bar plots. Bars are color-coded by module and reflect diverse functional themes including neurodegeneration, oxidative stress, inflammation, and metabolic regulation. “Enrichment” refers to a standard term in gene set analysis denoting the statistically significant overrepresentation of a defined gene set within a ranked list of genes, as computed by gene set enrichment analysis, GSEA.)

In more detail, the observation in the six modules were as follows: (see Figure 3A–D for corresponding temporal trajectories, GO enrichments, and KEGG pathways)

#### Module M1 (SCI)

Module M1 exhibited significant downregulation immediately following SCI, with gene expression sharply suppressed within 30 minutes and remaining low through 28 days. This rapid decline indicated immediate neuronal dysfunction due to conduction block, axonal injury, and subsequent neuronal death. Persistent suppression aligned with long-term neuronal and myelin loss, reflecting irreversible neural circuitry damage.

Module M1 was strongly associated with synaptic transmission, cognition, and memory functions, as evidenced by highly significant GO terms related to synaptic organization, vesicle transport, and exocytosis. KEGG pathways enriched included those linked to neurodegenerative diseases and calcium signaling, highlighting neuronal impairment and altered metabolic states post-injury. Notably, this module shows minimal internal ECM axis activation, contrasting with the concurrent HA, Provisional, and Protease engagement observed in other modules.

#### Module M2 (SCI)

Module M2 is downregulated from the onset of SCI, initially reflecting rapid oligodendrocyte gene suppression due to ischemic shock and metabolic stress. Gene expression remains significantly reduced at day 28, indicating prolonged demyelination and incomplete remyelination.

This module is notably enriched for mitochondrial functions, including aerobic respiration, oxidative phosphorylation, and reactive oxygen species (ROS) handling pathways. The enrichment patterns pointed to a metabolic crisis (Figure 3 C&D) in neurons and oligodendrocytes attempting to adapt to sustained oxidative stress and high energy demands post-injury. ECM signatures within this module remain weak; however, the persistent neuronal suppression coincides with heightened PNN-CSPG accumulation and fibrosis in other modules, potentially limiting functional recovery.

#### Module M3 (SCI)

Module M3 is robustly and persistently upregulated beginning at one day post-SCI, signifying sustained microglial and macrophage activation. Gene expression peaks by day seven and remains high through chronic injury phases, reflecting ongoing inflammation and innate immune activation typical in SCI lesions. This module is primarily associated with immune processes such as antigen processing, phagocytosis, and presentation via MHC class II, the major histocompatibility complex, a family of cell-surface proteins critical for adaptive immune recognition,, indicating microglial transformation into antigen-presenting and phagocytic cells critical for debris clearance. Within the ECM-axis framework, Protease/regulator genes (MMP2, MMP9, ADAMTS4/5, TIMPs) are prominent, while HA turnover and Basement-membrane terms co-enrich and PNN-CSPG components begin to accumulate.

#### Module M4 (SCI)

Module M4 in SCI exhibits moderate expression changes, with subtle increases at later stages (7–28 days), indicating its involvement in long-term extracellular matrix (ECM) remodeling and cell survival processes. KEGG pathway analysis reveals enrichment in PI3K-Akt, focal adhesion, relaxin, and TNF signaling, all critical for cell survival, migration, and ECM interactions. PI3K-Akt signaling promotes cell survival and proliferation, often activated downstream of ECM receptor signaling. Focal adhesion pathways suggest integrins binding ECM proteins such as collagen and fibronectin, initiating intracellular cascades important for cellular migration and structural organization. Axis mapping: Provisional matrix persists (FN1/TNC/integrins), Basement-membrane remodeling is extended, and PNN-CSPG and Crosslinking/fibrosis terms increase, marking transition toward a consolidated scar. Notably, SCI maintains ECM remodeling module activity substantially longer than BI.

#### Module M5 (SCI)

Module M5 shows rapid and intense upregulation immediately after SCI, peaking at around 4 hours and remaining significantly elevated at 1 day. However, its activity declines sharply by 3 days, falling below baseline at 7 days, and becomes suppressed by 28 days. This module primarily represents the acute inflammatory "cytokine storm" phase characteristic of SCI. Critical genes within this module, such as CCL2, CCL3, IL-6, and immediate early transcription factors like c-Fos, reflect rapid activation of innate immunity pathways immediately after injury.

Gene Ontology (GO) enrichment strongly emphasizes processes like apoptosis regulation, innate immune activation, inflammatory response, leukocyte migration, and chemotaxis, highlighting Module M5’s core role in orchestrating early post-injury inflammation. KEGG pathways notably include cytokine-cytokine receptor interaction, NF-κB signaling, Tlr signaling, and IL-17 signaling, reinforcing this module’s function in acute immune response regulation. In ECM-axis terms, Hyaluronan and Provisional matrix components dominate the acute phase, while concurrent Protease activity establishes the substrate for later ECM turnover.

#### Module M6 (SCI)

Module M6 emerges significantly at 3–7 days post-SCI, remaining active into chronic phases beyond 28 days. Initially inactive at acute stages (4h–1d), this module is progressively upregulated as monocyte-derived macrophages infiltrate the lesion, and microglia shift towards a phagocytic phenotype. By day 7, expression is robust, with genes like ApoE and Trem2 markedly elevated, reaching peak activity by day 28. APOE was particularly noted for its consistent high expression in chronic SCI macrophages and microglia.

Gene ontology analyses highlight significant involvement in lipid metabolism, catabolic processes, and immune regulation. Key biological pathways include lipid catabolism, lysosomal activity, fatty acid breakdown, and glycan degradation. Within the ECM-axis framework, PNN-CSPG signatures strengthen, crosslinking/fibrosis components (Lox/Loxl2/TGM2; COL1/3) become dominant and persistent, and basement-membrane terms remain detectable, indicating prolonged neurovascular remodeling.

### Cross-Validation with Neural-Implant Transcriptomics

The discovery-phase analyses established HA as the top-ranked ECM axis in BI/SCI models. To test whether these findings, particularly the HA-DAMP signaling mechanism, translate to neural implant contexts, we analyzed an independent transcriptomic dataset from tissue adjacent to chronically implanted flexible polyimide thin-film probes.

Although our primary analysis focused on BI and SCI, the molecular signatures identified, particularly those associated with inflammation, oxidative stress, and ECM remodeling, exhibit substantial convergence with transcriptional programs reported in intracortical neural-implant models. Multiple groups have shown parallel temporal trajectories of immune activation and glial remodeling following microelectrode insertion and chronic implantation, reinforcing the broader relevance of injury-associated findings of this work. However, the present study moves beyond these descriptive parallels by identifying specific co-expression modules and their constituent ECM axes, providing a mechanistic framework that links immune activation to defined matrix remodeling programs.

In a recent systems-level study, Huff et al.(Huff et al. 2025) performed a time-resolved comparison of craniotomy (CO), stab-wound (SW), and implanted (Im) conditions at 2, 8, and 16 weeks. Their analysis revealed sustained upregulation of Hmox1, Il6, Ccl2, Cd14, Cd44, Socs3, Tlr2, Tlr4, Gfap, and Vim in the implant group, whereas expression in CO and SW controls declined toward baseline. These findings confirm persistent oxidative stress signaling, monocyte recruitment, and astrocytic activation at the device–tissue interface, molecular processes that closely parallel the acute-phase and subchronic responses identified in BI and SCI networks.

Complementary short-term data from Joseph et al.(Joseph et al. 2021) capture the acute response within hours after probe insertion. Early enrichment of Mmp12, Mmp9, Mmp14, and Mmp19 marked rapid ECM disorganization and neutrophil degranulation, while reactive-astrocyte modules showed increased Cd14, Cd44, Lcn2, Gfap, Vim, Timp1, Serping1, and Tgm1 expression, driven by JAK-STAT and TNF signaling. Subsequent imaging confirmed high GFAP and IBA1 expression adjacent to the implant, followed by partial resolution of astrocyte reactivity and restoration of neuronal density by 18 weeks.

Complementary work by Bennett et al.(Bennett et al. 2021) and Song et al.(Song et al. 2022) confirmed that cortical penetration and probe presence elicit robust activation of innate-immune pathways, Il6, Cxcl1, Cxcl10, Socs3, and Tlr2/4, together with classical astrocytic and matrix-remodeling markers (Gfap, Vim, Fn1, Mmp2, Mmp9). These signatures closely mirror the inflammatory and reparative modules characteristic of traumatic-CNS-injury paradigms.

These temporal patterns align precisely with our BI/SCI-derived co-expression modules encompassing HMOX1-, IL6-, and Tlr-mediated inflammation and Cd44-linked ECM remodeling. The reproducibility of these gene networks across mechanical injury and implant contexts suggests that they represent core neuroimmune–matrix programs rather than model-specific artifacts. Nevertheless, direct experimental validation in controlled neural-implant systems remains necessary to establish the temporal hierarchy and mechanistic coupling between oxidative stress, cytokine release, and matrix feedback. Such cross-evaluation would also clarify how hyaluronan (HA) fragmentation modulate immune tone and repair dynamics at the electrode–tissue interface, a topic currently characterized by heterogeneous and inconsistently applied definitions. To quantify this convergence, we evaluated whether BI/SCI-derived modules show significant overlap with published implant gene signatures (Figure 4).

**Figure 4.**
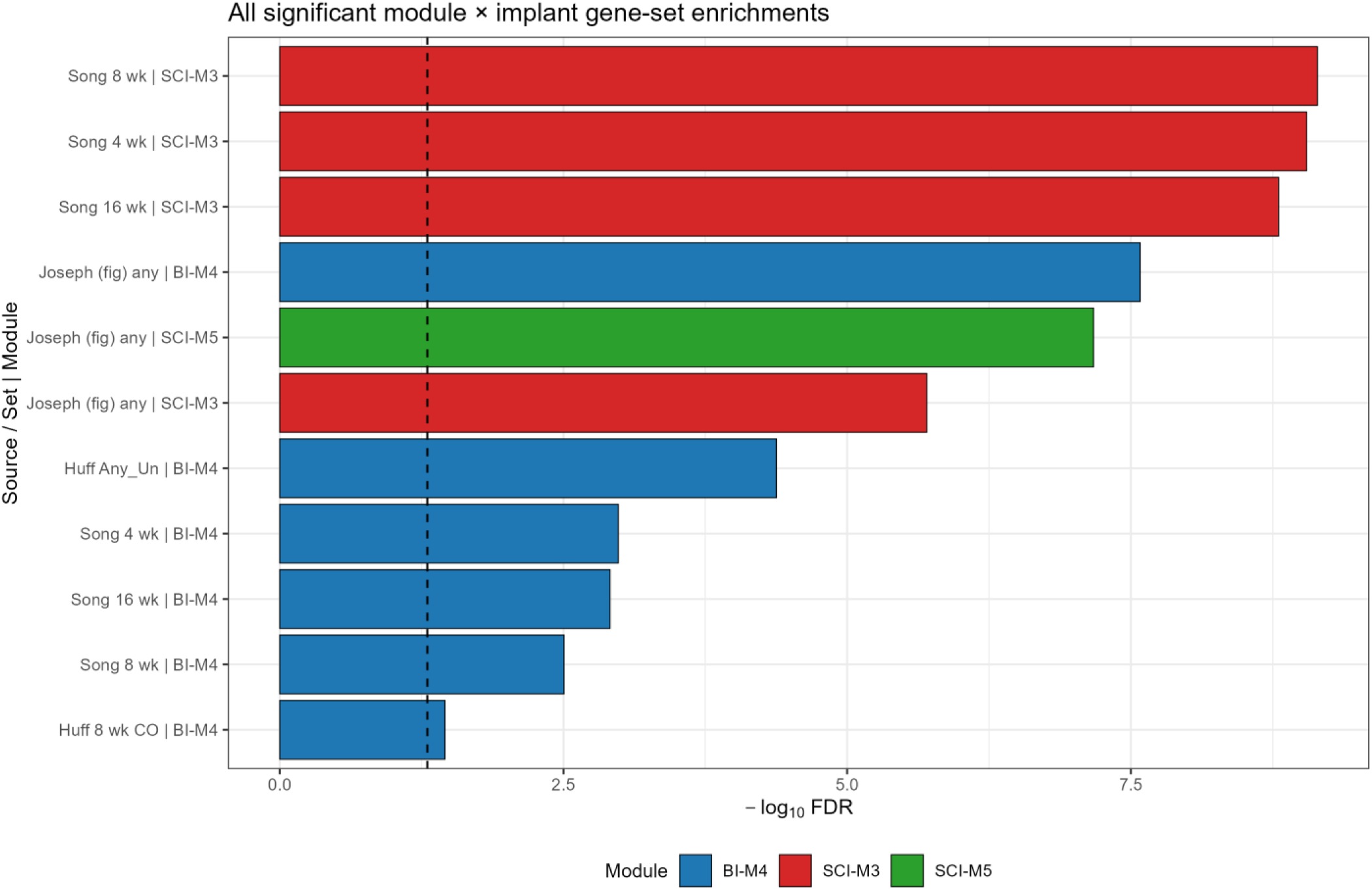
All significant module × implant gene-set enrichments (−log₁₀ FDR). Each bar represents a statistically significant overlap (Fisher’s exact test, FDR < 0.05) between a WGCNA co-expression module identified in the BI/SCI discovery analysis and an independently published implant-response gene set; taller bars indicate stronger enrichment. Bars show overlaps for Song (4/8/16 wk; panel-restricted background), Huff et al. 2025 (8 wk CO and time-collapsed Any_Un; panel-restricted), and a conservative figure-derived set from Joseph et al. 2021 (genome-wide array background). Colors indicate WGCNA modules (BI-M4 blue, SCI-M3 red, SCI-M5 green).

BI/SCI modules showed statistically significant overlap with gene-expression signatures from intracortical microelectrode models. Using Song et al. 2022 and a panel-restricted background, BI–M4 was enriched at 4, 8, and 16 weeks (OR = 5.51–7.27; FDR = 1.0×10⁻³–1.2×10⁻³), while SCI–M3 showed very strong enrichment at all time points (OR = 15.06–25.97; FDR = 8.9×10⁻¹⁰–1.6×10⁻⁹). With Huff et al. 2025 (2/8/16 wk; Un/CO/SW/Im), directionally consistent results were obtained, for example, BI–M4 at 8-week craniotomy-only yielded OR ≈ 6.16 (FDR ≈ 0.035), and time-collapsed union sets increased power (e.g., BI–M4 vs Any_Un: OR = 6.62, FDR = 4.2×10⁻⁵). A conservative, figure-derived implant panel (a targeted gene panel used in neural implant validation study from Joseph et al.) from Joseph et al. 2021 further supported convergence, with significant enrichment for BI–M4 (OR = 21.7; FDR = 2.6×10⁻⁸), SCI–M5 (OR = 34.7; FDR = 6.7×10⁻⁸), and SCI–M3 (OR = 13.9; FDR = 2.0×10⁻⁶).

Because the implant datasets employed targeted gene panels or broad hybridization arrays rather than whole-transcriptome sequencing, enrichment analyses used platform-matched background universes, panel-restricted (limited to the ∼770 neuroinflammation-associated genes on the NanoString nCounter® panel used by Song et al. and Huff et al.) or array-restricted (limited to the ∼20,000 genes represented on the hybridization array used by Joseph et al.), and these findings were accordingly treated as supportive but not primary. A stronger test of ECM-axis convergence came from a whole-transcriptome implant dataset, which, unlike the panel-restricted comparisons above, captured the full breadth of transcriptional programs activated around chronically implanted flexible polyimide probes.

#### HA-DAMP Pathway Ranks Among Top Implant-Activated Programs

To directly test whether the BI/SCI-derived ECM axis framework generalizes to neural implant contexts, we analyzed transcriptomic data from rat cortical tissue adjacent to flexible polyimide probes across five post-implantation timepoints (4h, 7d, 14d, 28d, and 126d; n = 63 samples). Gene set enrichment analysis revealed that HA-DAMP Signaling (*Cd44*, *Tlr2/4*, *Cd14*, and NFkB) ranked among the most consistently activated pathways, with a mean NES of 2.05 and significant enrichment at 4 of 5 timepoints (Figure 5a). This placed HA-DAMP signaling alongside literature-validated implant signatures (Huff et al., NES = 2.24; Joseph et al., NES = 2.17) and the Provisional Matrix axis (NES = 2.06), and above other canonical ECM programs including Proteases/Regulators and Crosslinking/Fibrosis. Notably, broader HA metabolism gene sets showed substantially weaker enrichment (NES = 0.66–1.50, 0–2/5 timepoints significant), suggesting that the implant transcriptional response is more consistent with HA-DAMP receptor activation than with homeostatic HA turnover.

**Figure 5.**
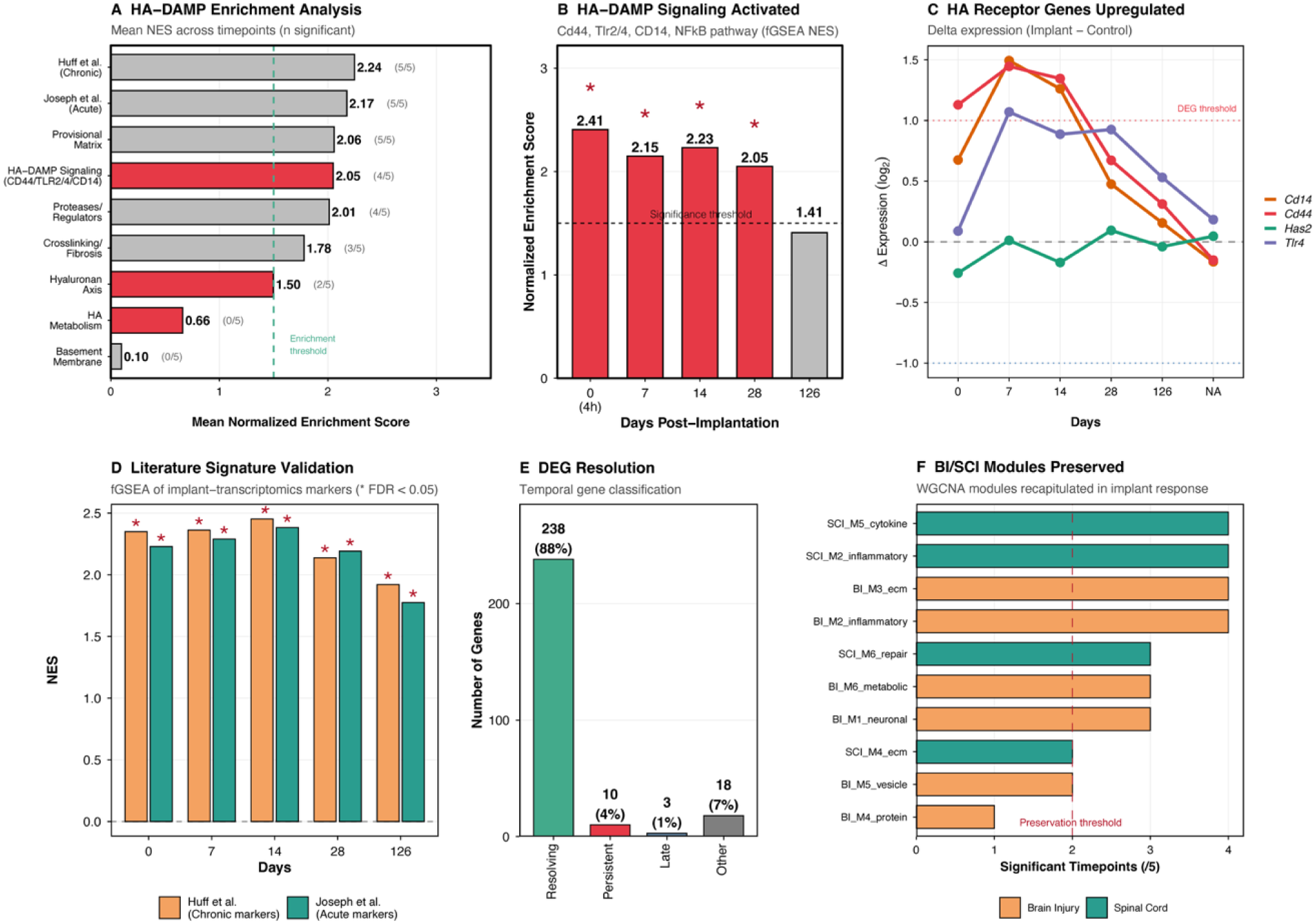
Validation of Hyaluronan-DAMP signaling as central orchestrator of neural implant ECM response. Transcriptomic analysis of rat cortical tissue adjacent to flexible polyimide neural probes (n = 63 samples across 5 timepoints: 4h, 7d, 14d, 28d, 126d) validates the ECM axis framework derived from brain injury (BI) and spinal cord injury (SCI) datasets. (A) Pathway enrichment ranking by mean normalized enrichment score (NES) across all timepoints (fGSEA). HA-DAMP Signaling (Cd44, Tlr2/4, Cd14, NF-κB targets; red bar) ranked 4th overall (NES = 2.05, 4/5 timepoints significant), comparable to literature-validated implant signatures and canonical ECM pathways. Broader HA metabolism and turnover gene sets showed weaker enrichment (NES = 0.66–1.50). Parentheses indicate timepoints with FDR < 0.05; dashed line marks enrichment threshold (NES = 1.5). (B) Temporal profile of HA-DAMP pathway enrichment. NES exceeded 2.0 at 4 of 5 timepoints (*FDR < 0.05), peaking at 4h (NES = 2.41) and remaining elevated through day 28 before declining below threshold by day 126 (NES = 1.41). (C) Individual gene trajectories for HA pathway components. Receptor genes (Cd44, Cd14, Tlr4) showed coordinated upregulation peaking at day 7 (Δ = 1.0–1.5 log₂), while the synthesis gene Has2 remained unchanged. This dissociation indicates upregulation of HA fragment sensing machinery without compensatory HA production (see Supplementary Figure S1 for complete HA turnover gene analysis). (D) Cross-validation with published neural implant signatures. Gene sets from Huff et al. (chronic markers: Hmox1, Il6, Ccl2, Cd44, Gfap, Vim) and Joseph et al. (acute markers: Mmp9, Mmp12, Lcn2, Timp1) showed significant enrichment at all timepoints (*FDR < 0.05), confirming convergent transcriptional programs across implant studies. (E) Temporal classification of differentially expressed genes. Of 269 genes significant at ≥ 1 timepoint, 238 (88%) showed resolving patterns, 10 (4%) were persistently dysregulated, and 3 (1%) emerged only at late timepoints. NA, not classifiable (n = 18, 7%). (F) WGCNA module preservation analysis. Bars indicate number of timepoints (of 5) at which BI-and SCI-derived modules showed significant enrichment in implant data. Nine of ten modules exceeded the preservation threshold (≥2 timepoints; dashed line), with inflammatory (M2/M5) and ECM remodeling (M3/M4) modules most strongly preserved.

#### Temporal Dynamics and Receptor Upregulation

Temporally, the HA-DAMP pathway showed peak activation at 4 hours post-implantation (NES = 2.41), remained significantly elevated through day 28 (NES = 2.05), and declined below the enrichment threshold by day 126 (NES = 1.41; Figure 5b). This trajectory, acute activation with eventual resolution, is consistent with a transient inflammatory response to insertion trauma. Individual gene analysis confirmed this pattern. HA receptor genes showed coordinated upregulation peaking at day 7 (Δ = 1.0–1.5 log₂), whereas the synthesis gene *Has2* remained at baseline (Figure 5c). Extended analysis of all HA turnover genes revealed a striking dissociation: receptor genes were selectively upregulated while neither degradation enzymes (*Hyal1–3*, *Spam1*) nor synthesis enzymes (*Has1–3*) showed significant changes at any timepoint (Supplementary Figure S2). *Tlr2* and *Hmmr* also remained unchanged, suggesting *Tlr4*/*Cd44*/*Cd14* as the dominant HA-sensing axis. This expression pattern indicates that LMW-HA fragments driving the DAMP response were generated by initial insertion trauma rather than ongoing enzymatic degradation.

#### Cross-Validation, Biocompatibility Dynamics, and Module Preservation

Cross-validation with published neural implant transcriptomic signatures confirmed convergent biology across studies. Gene sets derived from Huff et al. (chronic markers) and Joseph et al. (acute markers) both showed significant enrichment at all five timepoints (Figure 5d), demonstrating that flexible polyimide probes elicit transcriptional programs consistent with the broader implant literature. Temporal classification of differentially expressed genes revealed favorable biocompatibility dynamics: of 269 DEGs identified across all timepoints, 238 (88%) showed resolving patterns, returning to baseline by day 126 (Figure 5e). Only 10 genes (4%) were persistently dysregulated, and 3 (1%) emerged exclusively at late timepoints. Finally, WGCNA module preservation analysis demonstrated that 9 of 10 BI/SCI-derived co-expression modules were recapitulated in implant tissue, exceeding the preservation threshold of significance at ≥2 timepoints (Figure 5f). The most strongly preserved modules were those associated with acute inflammation (SCI-M5, BI-M2) and ECM remodeling (BI-M3, SCI-M4), while protein synthesis modules showed weaker preservation.

Collectively, these cross-evaluation analyses establish three findings: (i) HA-DAMP receptor signaling ranks among the strongest transcriptional programs activated by neural implants, consistent with a central role in the implant tissue response; (ii) the response is characterized by receptor upregulation without corresponding changes in HA turnover enzymes, consistent with trauma-generated rather than enzymatically-produced HA fragments; and (iii) 90% of injury-derived co-expression modules are preserved in implant tissue, supporting the use of BI/SCI datasets as mechanistically relevant surrogates for neural interface biology.

## Discussion

Returning to the core objectives stated in the Introduction, firstly, the performed WGCNA analysis of BI and SCI transcriptomes identified six conserved ECM-associated gene modules, with hyaluronan (HA) metabolism and fragmentation dynamics emerging as the most consistently activated ECM axis across both injury types. Our analysis supports a model in which extracellular matrix (ECM) remodeling, and in particular, HA-associated signaling, is consistently linked to the immune response observed in both neural injury models. Furthermore, for the first time, we demonstrate that gene programs associated with HA dynamics and their underlying networks are engaged across both injury models (BI, SCI) and neural implant-induced injury, motivating closer examination of HA-associated pathways as a unifying framework. Together, these results support our initial hypothesis that implant-induced injury engages injury programs characteristic of traumatic brain injury.

A key finding from the module-level analysis is that BI and SCI engage the same six ECM axes but with markedly different temporal profiles. BI compresses the ECM response into a brief but coordinated sequence, marked by a rapid hyaluronan and provisional matrix surge, moderate protease activation, transient Basement-membrane repair, limited PNN-CSPG engagement, and minimal fibrosis. In contrast, SCI expands and prolongs this program, with sustained hyaluronan and provisional activity, extended basement-membrane remodeling, broader protease participation, pronounced PNN-CSPG accumulation, and persistent crosslinking/fibrosis, with components such as Lox, Loxl2, TGM2, COL1, and COL3 becoming dominant in chronic SCI phases, consistent with consolidation into a fibrotic, crosslinked matrix that persists well beyond the reparative phase observed in BI. This divergence in ECM trajectory has direct implications for neural implant biocompatibility.

Similar to the investigated brain and spinal cord injury paradigms, the introduction of a neural implant inevitably produces acute mechanical and vascular trauma, e.g. shearing delicate parenchyma, rupturing microvessels, and perturbing local homeostasis. Previous studies suggest that mechanical and oxidative insults can fragment extracellular-matrix components, including hyaluronan (HA). In particular, high-molecular-weight HA (HMW-HA) can be structurally degraded by mechanical shear, reactive oxygen species (ROS), and enzymatic activity, to lower-molecular-weight HA fragments (Termeer et al. 2002; Scheibner et al. 2006). As for now, in the context of implant-induced injury, HA metabolism and the extent to which mechanical, oxidative, and enzymatic factors may alter HA structure remain underexplored at the systems-transcriptional level.

Addressing the second aim of the study, our independent validation in neural implant datasets showed that the transcriptional response following device insertion is consistent with robust HA-DAMP signaling activation (NES = 2.05, 4/5 timepoints significant), with 9 out of 10 injury-derived modules preserved in the implant context.

Our co-expression network analyses revealed that Module M2 (BI) and Module M5 (SCI) show the strongest enrichment for innate immune activation and TLR signaling pathways. The immediate consequence of these activated modules is the release of pro-inflammatory cytokines (e.g., Il-1β, TNF-α) by macrophages and microglia, and recruitment of neutrophils, potentially creating a feed-forward loop of inflammatory damage. The indicated correlations are consistent with previous literature suggesting distinct biological roles for different HA molecular weights (Stern et al. 2006), where, HMW-HA has been shown to foster environments supportive of proliferation, tissue stability, and regenerative healing processes, whereas LMW-HA fragments are well-established as pro-inflammatory signals that promote angiogenesis, cell migration, and differentiation associated with active inflammation and tissue remodeling (Jiang et al. 2007; Stern et al. 2007).

While our transcriptomic analysis identified gene modules linked with HA-associated biological pathways, these correlations do not yet establish direct causation with regards to HA molecular weight. Nevertheless, the convergent evidence suggest that HMW-HA likely conveys a tissue-stabilizing, immunomodulatory signal indicative of healthy ECM conditions, while LMW-HA serves as a “danger signal,” indicative of ECM damage and inflammation, concepts consistent with our hypothesis framework on ECM-immune integration.

In line with the literature, both BI and SCI demonstrate a biphasic response characterized by initial HA breakdown followed by subsequent HA synthesis (Struve et al. 2005; Xing et al. 2014). This dynamic is closely linked to progression of inflammation and repair mechanisms (Sherman et al. 2015; Srivastava et al. 2020). In the acute phase post-injury, both BI and SCI induce oxidative stress that degrades HMW-HA into LMW-HA (Soltés et al. 2006). This early oxidative and mechanical breakdown mirrors the acute phase observed around neural implants, dominated by innate inflammation and ROS-driven ECM remodeling (Ereifej et al. 2018; Bennett et al. 2019; Kozai et al. 2015). These LMW-HA fragments universally act as danger signals, activating innate immune pathways through Toll-like receptors (Tlr s), particularly via the MyD88/NF-κB axis, inducing pro-inflammatory cytokines such as IL-1β, IL-6, and TNF-α (Termeer et al. 2002; Scheibner et al. 2006; Jiang et al. 2005). This acute inflammatory response is common to both injury types, featuring significant leukocyte infiltration and a marked increase in reactive oxygen species (ROS), directly contributing to secondary tissue damage (Byrnes et al. 2007).

Subsequent to this inflammatory surge, glial cells in both BI and SCI shift towards increased HA synthesis, primarily through hyaluronan synthase 2 (HAS2), resulting in re-accumulation of HMW-HA in the ECM (Xing et al. 2014; Al’Qteishat et al. 2006). This reformed HA scaffold constitutes an integral part of the nascent glial scar, providing structural support and serving as a platform for binding other ECM molecules, notably proteoglycans such as neurocan and versican (Asher et al. 2002; Matsumoto et al. 2003). The receptor Cd44, upregulated on astrocytes and microglia post-injury, mediates these effects by internalizing and clearing HA fragments and transducing signals from HMW-HA to reduce inflammation and facilitate repair (Moon et al. 2004; Knudson et al. 2002).

Thus, despite anatomical differences, BI and SCI follow a conserved sequence of ECM remodeling and immune modulation: HA degradation leading to inflammation via LMW-HA/Tlr signaling, followed by HA synthesis and scar formation mediated through HMW-HA/Cd44 interactions (Struve et al. 2005; Al’Qteishat et al. 2006; Termeer et al. 2002; Scheibner et al. 2006; Moon et al. 2004).

System-level WGCNA module analyses underscore these shared molecular mechanisms. For instance, innate immune and inflammatory modules (BI M2, SCI M5) both highlight overlapping cytokine signaling and Tlr pathways, affirming the conserved inflammatory role of LMW-HA fragments. Similarly, modules enriched for oxidative stress (BI M1/M6, SCI M2) underline ROS-driven HA degradation processes shared across both injuries. Furthermore, modules associated with phagocytosis and antigen presentation (BI M3, SCI M3) indicate a common necessity for debris clearance and inflammation resolution. Effective clearance via Cd44-positive phagocytes correlates strongly with improved outcomes, as residual HA debris perpetuates chronic inflammation. Modules linked to ECM remodeling and scar formation (BI M4, SCI M4) reflect the vigorous astrocytic activation, translational machinery engagement, and survival signaling pathways, highlighting their roles in synthesizing ECM components including HA. Despite these shared features, marked differences in HA’s role and persistence are apparent between BI and SCI.

The contrast between BI and SCI ECM responses is most apparent in fibrotic scar formation: SCI develops a more pronounced and persistent scar, as reflected by greater ECM and focal adhesion pathway enrichment in Module M4 and consistent with the dense, lasting fibrotic scar described after spinal cord injury (Bradbury und Burnside 2019). Spinal lesions typically exhibit a dense HA- and CSPG-rich perilesional boundary that significantly inhibits axon regeneration, whereas BI permits HA to diminish over time as tissue partially regenerates or cavities form, reflecting the brain’s greater plasticity. These differences are compounded by distinct cellular sources of HA: in the brain, meningeal cells and resident astrocytes are the primary HA producers post-injury (Sherman und Back 2008), while SCI recruits additional perivascular fibroblasts that deposit collagen and modulate HA metabolism through hyaluronidase secretion or 4-methylumbelliferone-sensitive pathways (Soderblom et al. 2013; Nagy et al. 2015). Consequently, SCI modules emphasize fibrosis-related genes such as collagens and periostin, whereas BI modules are weighted toward neuron–glia interactions. The two injury types also diverge in adaptive immune engagement: chronic SCI involves sustained B and T lymphocyte infiltration, potentially driven by HA fragments enhancing self-antigen presentation via microglial activation (Jones et al. 2002; Ankeny et al. 2009), while BI resolves inflammation more rapidly with less prominent adaptive immunity. This is evidenced by persistent HA-binding protein staining weeks after SCI, whereas HA-related changes are transient in brain injuries (Passaro et al. 2021). These anatomical and immunological differences between BI and SCI are reflected in the distinct module compositions and temporal profiles captured by the ECM-axis framework.

Even though the role of HA in tissue remodeling is not new and experimental targeting of HA in BI and SCI is reported in literature (Štepánková et al. 2023, Zheng et al. 2022, Washington et al. 2020), the precise underlying mechanisms, its predominant role among ECM signaling axes, and its specific relevance in the context of implant-induced injury have remained insufficiently understood.

Whether HA-associated signaling also influences adaptive immune priming remains speculative, though published studies suggest that HA molecular weight can bias immune tolerance versus activation (Johnson et al. 2018; Imani et al. 2021). In our data, antigen-processing modules (MHC class II–related genes) co-activate with LMW-HA–linked inflammatory signatures in both BI and SCI, but direct causal links between implant-induced HA fragmentation and adaptive immunity remain to be demonstrated.

Our findings offer new perspectives advancing biocompatibility measures in the context of implantable neural interfaces. Building on the concept of extracellular matrix (ECM) remodeling during neural implant–induced injury and the central role of hyaluronan (HA) fragmentation, the trauma-generated mechanism identified in this study carries direct implications for neural implant design. If LMW-HA fragments arise from mechanical and oxidative insult rather than sustained enzymatic activity, then design strategies should prioritize minimizing insertion trauma and preserving ECM integrity at the device-tissue interface.

This implies that an ideally biocompatible neural interface could be designed to minimize mechanical and biochemical drivers of HA degradation. In the context of intracortical micro electrodes arrays, HA degradation can arise from factors like, mechanical shearing during insertion and persisting micromotion (chronic), electrochemical reactions at the electrode surface that generate reactive oxygen species (ROS), and biological factors (Table 3). Thus, the aim should be to preserve high-molecular-weight HA (HMW-HA) within the peri-implant matrix, thereby maintaining a regenerative and anti-inflammatory microenvironment at the device–tissue interface. In other word, a biocompatible neural implant should neither degrade HA beyond what the tissue can naturally recover from, nor interfere with endogenous repair mechanisms.

**Table 3.**
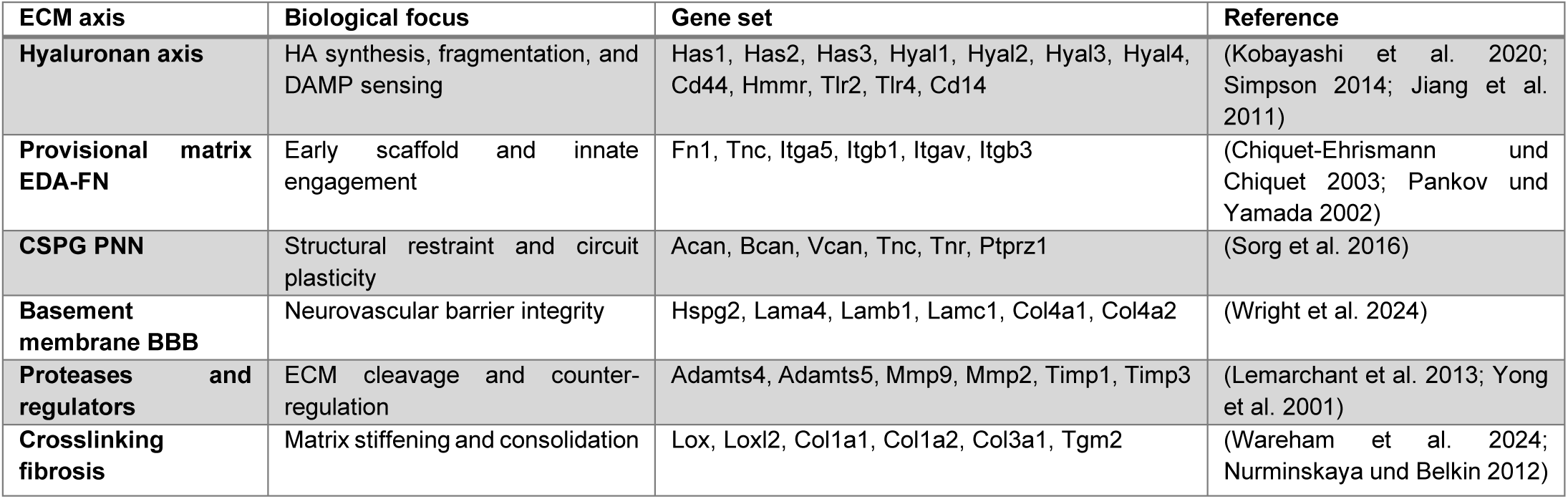
Extracellular matrix axes and curated gene sets.

By reducing micromotion, minimizing electrochemical by-products, and avoiding conditions that promote excessive ROS and protease activity, implant designs may help maintain a more quiescent, repair-supportive environment and lower the risk of chronic inflammatory scarring at the device–tissue interface. Electrochemical factors including reactive oxygen species (ROS) and pH shifts generated by faradaic charge injection, can fragment HA glycosidic linkages. Electrode materials and stimulation paradigms that favour capacitive coupling over faradaic reactions can substantially reduce HA fragmentation (Shepherd et al. 2021) when applied correctly, and HA-functionalized coatings have been shown to attenuate astrogliosis in vivo (Lee et al. 2015). Mechanical factors indued by shear stress from micromotion between rigid implants and compliant brain tissue, which physically tears HMW-HA; reducing bending rigidity through thin, filamentary geometries can minimize tissue-device mechanical mismatch even when the bulk modulus of the material remains high (Chernos et al. 2017; Vomero et al. 2020). Biological factors include enzymatic degradation by hyaluronidases, which are transcriptionally upregulated following CNS insults and actively digest HMW-HA into pro-inflammatory low-molecular-weight fragments within lesions (Sherman et al. 2015). Secondary biological cascades include sustained pH depression from cellular necrosis and continued ROS amplification by recruited phagocytes, creating a feed-forward loop of HA fragmentation. These biological pathways may be mitigated through anti-inflammatory coatings, ROS scavengers, or exogenous HMW-HA delivery (Jensen et al. 2020; Zheng et al. 2022; Isa et al. 2015).

By addressing electrochemical, mechanical, and biological degradation routes simultaneously, implant designs can avoid overwhelming the ECM’s regenerative capacity and maintain HA predominantly in its high-molecular-weight, anti-inflammatory form. An important next step is to define the threshold beyond which reparative high-molecular-weight HA turnover shifts toward sustained fragmentation and chronic inflammation. Quantifying this “ECM regenerative threshold” could be pursued through longitudinal liquid biopsy approaches, enabling minimally invasive monitoring of whether the implant–tissue interface remains within a reparative regime.

The insights from this work position hyaluronan (HA) not merely as a passive structural ECM component, but as a dynamic molecular integrator linking mechanical injury, innate inflammation, and longer-term immune remodeling. By revealing that HA-associated transcriptional programs are consistently engaged across brain injury, spinal cord injury, and also neural implant contexts, our analysis highlights HA-driven regulation as a previously underappreciated axis of biocompatibility. This framework suggests that HA-associated dynamics influence not only acute inflammatory responses, but also the trajectory toward resolution or chronic immune activation. Given that HA fragments and HA-modifying enzymes are detectable in cerebrospinal fluid and, under certain conditions, in peripheral circulation, we propose that HA-associated signatures warrant further investigation as candidate, non-invasive readout in the context of neural injury in general and tissue–implant interactions in particular.

### Scope and Limitations

Our inputs synthesize brain and spinal cord injury datasets, with implications for neural implants rather than implant-exclusive omics, so generalization to intracortical interfaces remains a hypothesis to be validated prospectively. The analysis relies on BI and SCI microarrays; transcript levels are incomplete proxies for protein activity and matrix biophysics. BI sampling covers only 7 days, whereas SCI extends to 28 days. This temporal mismatch means that the cross-model comparison primarily captures acute-to-subacute dynamics; extending BI datasets to later timepoints would be needed to confirm whether the converging ECM signatures observed here persist into the chronic phase. Future studies are needed to explicitly validate the causal mechanisms and confirm this molecular-weight-dependent functional dichotomy of HA. The validation dataset provides independent confirmation but is limited to a single probe type (flexible polyimide) in the rat cortex; broader generalization across implant materials and anatomical targets remains to be established.

## Conclusion

This study identifies hyaluronan (HA) metabolism as a key component of extracellular matrix (ECM) remodeling following neural injury. By integrating gene-expression and co-expression analyses across brain and spinal cord injury (BI and SCI) datasets, we demonstrate that HA-associated transcriptional programs are consistently engaged across injury contexts and closely associated with activation of innate immune pathways through Cd44 and Toll-like receptor signaling, inflammation resolution and tissue stabilization. These results underscore that HA ad presumably its molecular-weight dynamics govern the balance between inflammation and repair, providing a molecular framework for understanding how neural tissue responds to mechanical and biochemical stress.

Viewed from a biocompatibility perspective, these findings position HA not merely as a passive component of the ECM, but as a defining axis linking mechanical injury, innate inflammation, and longer-term immune remodeling in the injured central nervous system. Consistent with this, independent validation using neural implant transcriptomic data indicated that HA-DAMP signaling ranked among the strongest activated programs (NES = 2.05), and 90% of injury-derived co-expression modules showed significant preservation in implant tissue.

The transcriptional programs and ECM pathways we describe overlap with those previously reported around implanted electrodes, but our results, for the first time, position HA related pathways in the central role. By demonstrating preservation of HA-associated co-expression modules and strong enrichment of HA-DAMP signaling in implant tissue, this study provides molecular evidence that neural implants engage injury programs characteristic of traumatic brain injury rather than a conventional foreign-body response. As HA fragments and HA-modifying enzymes are detectable in cerebrospinal fluid and, under certain conditions, in peripheral circulation, HA dynamics may provide a minimally invasive readout of tissue–implant interactions. Such longitudinal monitoring of HA molecular-weight distributions, alongside targeted modulation of HA degradation or preservation, represents a promising strategy to gain insights on tissue remodeling processes following traumatic brain injury and neural implantation, an area that to date remains largely unexplored.

## Materials and Methods

### Dataset Selection for BI and SCI

To investigate gene expression changes associated with neural injuries and their potential links to ECM remodeling and immune modulation, two types of injuries were studied: brain injury and spinal cord injury, from publicly available microarray datasets from the National Center for Biotechnology Information (NCBI) Gene Expression Omnibus (GEO) repository (https://www.ncbi.nlm.nih.gov/geo/; accession numbers GSE35338 and GSE5296) (Edgar et al. 2002; Barrett et al. 2013). Two datasets, GSE35338 and GSE5296, were chosen based on their coverage of time-dependent neural injury responses relevant to astrogliosis:

1. Dataset GSE35338: This dataset contains gene expression profiles from mice subjected to middle cerebral artery occlusion (MCAO), a model for ischemic brain injury. Samples were harvested at 1, 3, and 7 days post-MCAO, alongside sham-operated controls. The dataset also includes expression profiles following lipopolysaccharide (LPS) injections, with samples taken 1 day post-injection to replicate inflammatory conditions.
2. Dataset GSE5296: This dataset captures gene expression profiles from mice after traumatic spinal cord injury (SCI). Samples were collected at multiple time points post-injury, including 0.5 hours, 4 hours, 1 day, 3 days, 7 days, and 28 days, compared to sham-operated controls at time zero.

These two series were selected because they (i) span acute–subacute phases in complementary Central Nervous System (CNS) injury contexts (brain vs spinal cord), (ii) were generated on the same Affymetrix platform (GPL1261), and (iii) therefore permit a unified preprocessing and normalization strategy across timepoints and models. Although neither model directly recapitulates device implantation, both share key pathophysiological features with implant-induced injury, including blood–brain barrier disruption, acute inflammation, and reactive gliosis, making them informative proxies for the early tissue response to neural implants. To avoid confounding systemic inflammation, the LPS arm in GSE35338 was excluded from discovery analyses; only Middle Cerebral Artery Occlusion (MCAO) and matched sham animals were analyzed. These two datasets, sourced from Affymetrix microarray experiments, offer temporal snapshots of the post-injury environment, allowing for the identification of DEGs associated with astrogliosis and more broadly with neural injury responses.

### Gene Expression Analysis

#### 1- Data Processing and Normalization

Both GSE35338 and GSE5296 datasets were generated on the Affymetrix Mouse Genome 430 2.0 Array platform (GPL1261). Raw expression data and sample annotations were downloaded from the GEO. Data normalization and initial statistical analyses were performed using GEO2R, a web-based tool that utilizes the R packages GEOquery (for data retrieval and parsing) and limma (for linear model-based differential expression analysis with empirical Bayes moderation) (Barrett et al. 2013). Expression intensities were log2-transformed to stabilize variance across experimental conditions. Multiple testing correction was applied using the Benjamini-Hochberg (BH) procedure, generating q-values (BH adjusted p-values) to control the expected proportion of false positives among genes called significant (false discovery rate, FDR < 0.05) (Benjamini und Hochberg 1995).

#### 2- Identification of Differentially Expressed Genes (DEGs)

For each time point, pairwise comparisons were conducted between injury samples (MCAO or SCI) and the respective sham controls. Differentially expressed genes (DEGs) were defined as those with Benjamini–Hochberg adjusted p-value (FDR/q-value) < 0.05 and an absolute log2 fold change (|log2FC|) ≥ 1.0. Each DEG was annotated with its official gene symbol, gene name, log2 fold change (logFC) value, and q-value for downstream analyses. Here, the p-value refers to the raw test probability for each gene, while the q-value represents the p-value after FDR adjustment to account for multiple hypothesis testing.

### Protein–Protein Interaction (PPI) Network Construction

#### 1- PPI Analysis Using the STRING Database

All identified DEGs were uploaded to the Search Tool for the Retrieval of Interacting Genes/Proteins (STRING; version 11.5, Mus musculus) to uncover putative protein-protein interactions (Szklarczyk et al. 2021). We used the default medium confidence cutoff (score ≥ 0.400) to balance recall and precision when building hypothesis-generating networks, which provides a broad interaction landscape suitable for discovery-phase screening. This threshold yields a network that incorporates experimental data, curated databases, and computational inferences, allowing the identification of both well-established and putative interactions. To mitigate the inclusion of spurious interactions inherent to permissive thresholds, downstream hub-gene prioritization was restricted to DEGs meeting stringent significance and effect-size criteria (FDR < 0.05 and |log₂FC| ≥ 2) and ranked by degree centrality in CytoHubba, thereby focusing the analysis on high-confidence, biologically meaningful nodes.

### 2- Network Visualization and Hub Gene Identification

Network data were imported into Cytoscape (version 3.8.2) for visualization and topological analysis (Shannon et al. 2003). Nodes represented individual genes or proteins, and edges indicated interactions predicted by the STRING database. Hub-gene prioritization was restricted to differentially expressed genes (DEGs) that satisfied the global significance criteria (Benjamini–Hochberg adjusted p-value (FDR) < 0.05 and |log₂FC| ≥ 2). This more stringent effect-size threshold (|log₂FC| ≥ 2 versus ≥ 1 used for global DEG calling) was applied to focus hub-gene analysis on the most biologically pronounced expression changes; all resulting hub genes also satisfy the primary DEG criteria (|log₂FC| ≥ 1, FDR < 0.05). These DEGs were ranked using the CytoHubba plugin, which applies degree centrality and related network metrics, and genes with the highest degree of centrality were considered candidate hub regulators of neural injury responses.

### Weighted Gene Co-expression Network Analysis (WGCNA)

To capture coordinated transcriptional programs, we constructed weighted gene co-expression networks separately for the brain-injury (BI; GSE35338) and spinal-cord-injury (SCI; GSE5296) datasets, thereby preserving context-specific topology; cross-dataset comparisons were performed at the module and enrichment levels rather than via a single consensus network. Networks were built using biweight midcorrelation with a signed topological overlap matrix (TOM). The soft-threshold power was evaluated independently for each dataset (Supplementary Figure S1). For the SCI network (n = 60), power = 6 achieved satisfactory scale-free topology fit (signed R² = 0.85). For the smaller BI dataset (n = 21), scale-free fit at power = 6 was lower (signed R² = 0.50), reflecting the higher noise floor in correlation estimates from small samples (Ballouz et al. 2015) and the known sensitivity of power-law fitting to limited data (Clauset et al. 2009). A common power was retained to enable direct cross-dataset module comparison; the biological coherence of the resulting BI modules, including strong injury correlation (M4: r = +0.79, p = 1.7 × 10⁻⁵) and convergent ECM gene enrichment, supports the validity of this choice. Modules were identified by dynamic tree cutting (minimum module size = 30; deepSplit = 2). Modules were merged using a height threshold of mergeCutHeight = 0.20 and a reassign Threshold of 0.80. Module eigengenes were computed and carried forward for correlation, enrichment, and cross-context overlap analyses.

### Functional Annotation and Pathway Analysis

Biological significance of the DEGs and hub genes was assessed via the Database for Annotation, Visualization, and Integrated Discovery (DAVID) to reveal over-represented Gene Ontology (GO) terms, including cellular components, molecular functions, and biological processes (Da Huang et al. 2009; Sherman et al. 2022). Enriched pathways were further examined using the Kyoto Encyclopedia of Genes and Genomes (KEGG) resource (Kanehisa et al. 2017).

### Definition of extracellular matrix axes and gene sets

Six extracellular matrix axes were defined by adapting the Matrisome classification framework (Naba et al. 2012; Naba et al. 2016) to CNS-specific injury biology. Naba’s Matrisome defines two divisions comprising six categories: the core matrisome (collagens, ECM glycoproteins, proteoglycans) and matrisome-associated proteins (ECM-affiliated proteins, ECM regulators, secreted factors). We transformed these into functional axes relevant to neural tissue response: (1) hyaluronan turnover and sensing, derived from proteoglycans, capturing molecular weight-dependent signaling relevant to CNS injury; (2) provisional matrix, derived from ECM glycoproteins, capturing acute injury-response molecules; (3) perineuronal-net CSPGs, derived from proteoglycans, isolating the lectican family specific to CNS barrier function; (4) basement membrane components, derived from collagens and glycoproteins, representing neurovascular scaffolding; (5) proteases and regulators, directly transferred from ECM regulators; and (6) crosslinking or fibrosis programmes, derived from ECM-affiliated proteins, capturing scar-forming processes. The gene lists, summarised below, were established before any downstream analysis.

To establish explicit connections between injury-responsive genes and ECM-related processes, a literature review was performed to locate studies that link specific genes to the ECM Axis. Relevant findings from KEGG and GO analyses were compared with known ECM-related pathways to pinpoint overlaps and potential regulatory axes, thereby elucidating how changes in gene expression might influence ECM and consequent immune modulation in the injured CNS (The Gene Ontology resource: enriching a GOld mine 2021; Kanehisa et al. 2021; Gillespie et al. 2022).

### Dataset and Analysis for Neural Implant Induced Injury

Transcriptomic data were obtained from a published neural implant study examining tissue responses to flexible polyimide probes in female Sprague-Dawley rats (Joseph et al. 2021). The dataset comprised n = 63 samples from 6 animals per timepoint: 31 implant samples and 32 paired sham controls across five timepoints (Day 0/4h, Day 7, Day 14, Day 28, and Day 126 post-implantation). RNA was extracted from 1 mm radius tissue surrounding the implantation site and hybridized to Affymetrix Clariom S Rat arrays (ThermoFisher). The paired experimental design enabled within-subject comparisons that control for inter-animal variability.

Raw CEL files were processed using the oligo package in R. Robust Multi-array Average (RMA) normalization was applied for background correction, quantile normalization, and median-polish summarization. Probe-to-gene mapping was performed using the clariomsrattranscriptcluster.db annotation package, with multi-mapping probes collapsed by selecting the probe with the highest mean expression per gene symbol. Differential gene expression between implant and control conditions was assessed at each timepoint using the limma package with a paired design accounting for animal-level correlation. Genes with |log₂ fold change| ≥ 1.0, and Benjamini-Hochberg adjusted p-value < 0.05 were considered differentially expressed genes (DEGs).

Pathway-level activity scores were calculated using single-sample Gene Set Enrichment Analysis (ssGSEA) via the GSVA package. Pre-ranked gene set enrichment analysis was performed using the fgsea package with 10,000 permutations. Genes were ranked by the signed -log₁₀(p-value) × sign(logFC) statistic. In addition to ECM axes, literature-derived gene signatures from Huff et al. and Joseph et al. were included. Genes significant (FDR < 0.05, |log₂FC| ≥ 1) at two or more timepoints were classified as Resolving (significant at early timepoints but not Day 126), Persistent (significant at both early and late timepoints), or Late-emerging (not significant at Day 0–14 but significant at Day 28 or 126).

## Code Availability

All analysis scripts for both discovery (BI/SCI) and validation (neural implant) phases are available at https://github.com/asharbatian/ECM_Axis_Neural_Injury_Analysis. Software packages and versions are detailed in the Supplementary Materials.

## Ethical Considerations

All data analyzed in this study originated from publicly accessible archives and did not require new animal or human experimentation. No additional institutional review board or animal research committee approval was necessary because the research focused exclusively on secondary analysis of existing microarray data.

## Acknowledgements

This work was partially supported by BrainLinks-BrainTools, funded by the Federal Ministry of Economics, Science and Arts of Baden-Württemberg within the sustainability program of the Excellence Initiative II, as well as by the BMBF (DiaQNOS, 1010 0665 01) and the DFG (MA 10605/1-1).

