## Supplementary Material for "In Search for Biomarkers Reflecting Neural Implant-Induced Tissue Response Dynamics"

*[1] Laboratory for Biomedical Microtechnology, Department of Microsystems Engineering (IMTEK), University of Freiburg, 79110 Freiburg, Germany [2] BrainLinks-BrainTools, Institute for Machine-Brain Interfacing Technology (IMBIT), University of Freiburg, 79110 Freiburg, Germany [3] Laboratory of NeuroEngineering, Department of Neurosurgery, Medical Center- University of Freiburg, 79108 Freiburg, Germany [4] Section for Neuroelectronic Systems, Department for Neurosurgery, University Medical Center Freiburg, 79108 Freiburg, Germany [5] Department of Stereotactic and Functional Neurosurgery, Medical Center - University of Freiburg, Freiburg, Germany, 79106 Freiburg, Germany; [6] Faculty of Medicine, University of Freiburg, 79110 Freiburg, Germany*

⨼ These authors contributed equally to this work

**Table S1.** Summary of datasets, platforms, models, time points, and sample sizes used in the BI and SCI analyses. *Abbreviations:* BI, brain injury; SCI, spinal cord injury; MCAO, middle cerebral artery occlusion.

| Dataset | Platform | Injury model | Time points (injury vs control) | n per time point |
|---|---|---|---|---|
| GSE35338 (BI) | Affymetrix Mouse Genome 430 2.0 (GPL1261) | MCAO (brain) | 1 d, 3 d, 7 d vs matched shams | 1 d: BI = 5, sham = 4; 3 d: BI = 3, sham = 3; 7 d: BI = 3, sham = 3 † |
| GSE5296 (SCI) | Affymetrix Mouse Genome 430 2.0 (GPL1261) | T9 contusion (spinal cord) | 0.5 h, 4 h, 1 d, 3 d, 7 d, 28 d vs uninjured control | Control = 6; each injured time point = 9 |

† LPS/Saline arms in GSE35338 were excluded to avoid systemic-inflammation confounding.

Supplementary Figure S1. Scale-free topology fit diagnostics for soft-threshold power selection in WGCNA.

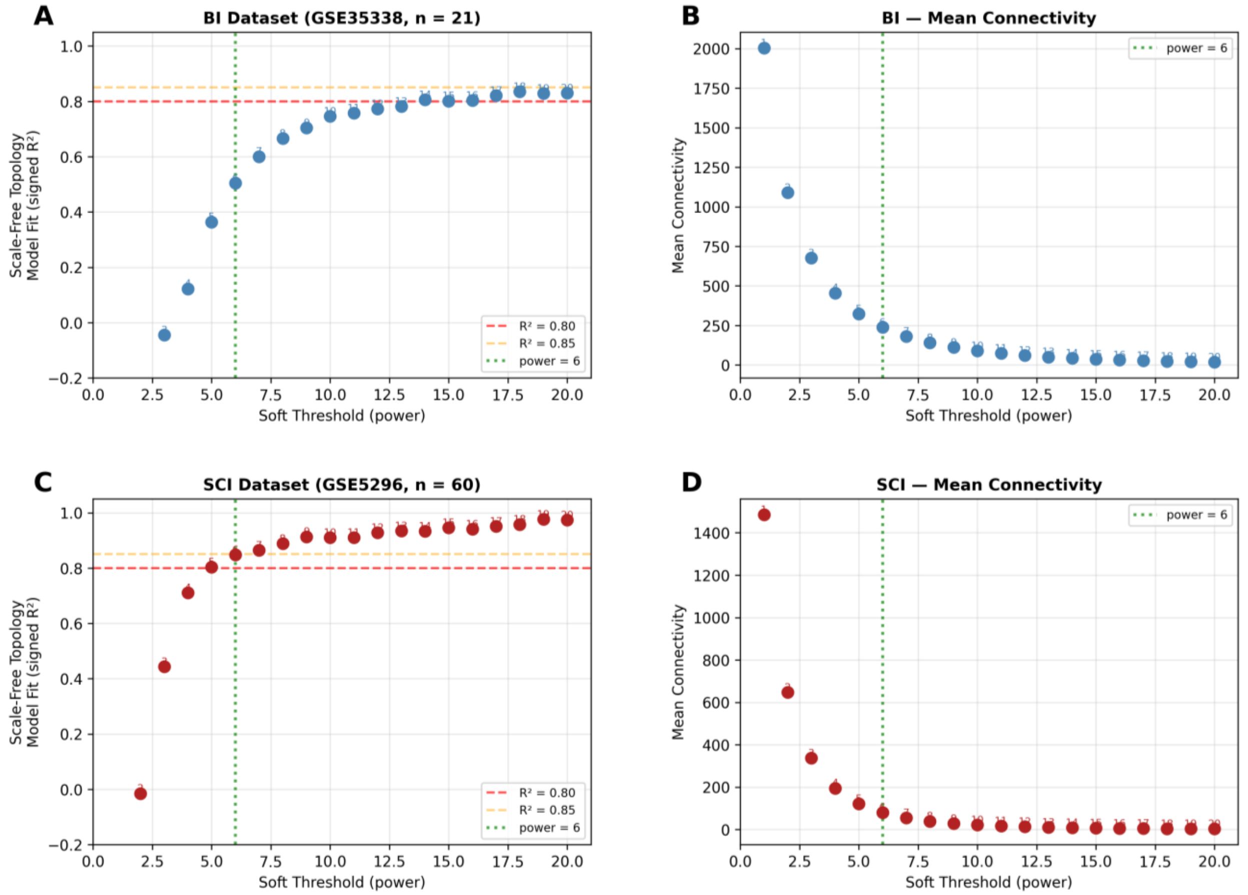


*Figure S1 Soft-threshold power was evaluated independently for BI (GSE35338, n = 21) and SCI (GSE5296, n = 60) using pickSoftThreshold (WGCNA, bicor correlation). (A, C) Signed $R^2$ versus candidate powers (1–20); dashed lines at $R^2 = 0.80$ and 0.85; green line marks selected power = 6. SCI achieved $R^2 = 0.85$ at power = 6. BI showed lower fit ($R^2 = 0.50$), expected for small sample sizes; biological validity is supported by strong module–injury correlations (M4: $r = +0.79$, $p = 1.7 \times 10^{-5}$). A common power was used for cross-dataset comparability. (B, D) Mean connectivity versus power.*

**Table S2.** Module-Trait Correlations for All WGCNA Modules with ECM Gene Mapping.

| Brain Injury (BI) - All 20 Modules (Sorted by Size) | | | | | | | | | |
|---|---|---|---|---|---|---|---|---|---|
| **Rank** | **Module Color** | **Genes** | **Injury r** | **p-value** | **Sig.** | **Top 6 (Size)** | **Trait (p<0.05)** | **ECM** | **ECM Gene List** |
| 1 | turquoise | 3283 | -0.074 | 0.748 | | Yes (M1) | No | 0 | - |
| 2 | blue | 1753 | +0.453 | 3.9e-02 | * | Yes (M2) | Yes | 7 | Tlr4, Itgb1, Lama4, Lamc1, Col4a1, Adamts4, Tgm2 |
| 3 | brown | 1668 | -0.123 | 0.596 | | Yes (M3) | No | 0 | - |
| 4 | yellow | 1637 | +0.794 | 1.7e-05 | *** | Yes (M4) | Yes | 14 | Has1, Has2, Cd44, Tlr2, Fn1, Tnc, Itga5, Vcan, Lamb1, Col4a2, Timp1, Lox, Loxl2, Itgb1 |
| 5 | green | 1568 | +0.010 | 0.966 | | Yes (M5) | No | 0 | - |
| 6 | red | 1174 | -0.239 | 0.296 | | Yes (M6) | No | 0 | - |

| | | | | | | | | | |
|---|---|---|---|---|---|---|---|---|---|
| 7 | black | 1046 | -0.653 | 1.3e-03 | ** | No | Yes | 2 | Bcan, Timp3 |
| 8 | pink | 787 | +0.387 | 0.083 | | No | No | 0 | - |
| 9 | magenta | 758 | -0.781 | 2.9e-05 | *** | No | Yes | 0 | - |
| 10 | purple | 735 | -0.640 | 1.8e-03 | ** | No | Yes | 0 | - |
| 11 | greenyellow | 639 | -0.324 | 0.152 | | No | No | 0 | - |
| 12 | tan | 556 | +0.317 | 0.162 | | No | No | 0 | - |
| 13 | salmon | 546 | -0.115 | 0.619 | | No | No | 0 | - |
| 14 | cyan | 386 | -0.258 | 0.260 | | No | No | 0 | - |
| 15 | midnightblue | 340 | -0.565 | 7.7e-03 | ** | No | Yes | 1 | Col1a1 |
| 16 | lightcyan | 335 | -0.370 | 0.099 | | No | No | 0 | - |
| 17 | lightgreen | 280 | +0.571 | 6.8e-03 | ** | No | Yes | 1 | Adamts5 |
| 18 | grey60 | 280 | -0.524 | 1.5e-02 | * | No | Yes | 1 | Ptprz1 |
| 19 | lightyellow | 244 | +0.565 | 7.6e-03 | ** | No | Yes | 0 | - |
| 20 | royalblue | 234 | +0.290 | 0.202 | | No | No | 0 | - |

**Spinal Cord Injury (SCI) - All 18 Modules (Sorted by Size)**

| Rank | Module Color | Genes | Injury r | p-value | Sig. | Top 6 (Size) | Trait (p<0.05) | ECM | ECM Gene List |
|---|---|---|---|---|---|---|---|---|---|
| 1 | turquoise | 1990 | -0.502 | 4.4e-05 | *** | Yes (M1) | Yes | 2 | Acan, Bcan |
| 2 | blue | 1687 | -0.337 | 8.6e-03 | ** | Yes (M2) | Yes | 0 | - |
| 3 | brown | 1552 | +0.251 | 0.053 | | Yes (M3) | No | 0 | - |
| 4 | yellow | 1192 | -0.178 | 0.174 | | Yes (M4) | No | 0 | - |
| 5 | green | 1160 | +0.303 | 1.8e-02 | * | Yes (M5) | Yes | 17 | Cd44, Tlr2, Tlr4, Cd14, Tnc, Itga5, Itgav, Lama4, Lamc1, Col4a1, Adamts5, Timp1, Timp3, Lox, Loxl2, Tgm2, Fn1 |
| 6 | red | 950 | +0.115 | 0.383 | | Yes (M6) | No | 0 | - |
| 7 | black | 691 | +0.292 | 2.4e-02 | * | No | Yes | 1 | Adamts4 |
| 8 | pink | 542 | -0.109 | 0.407 | | No | No | 0 | - |
| 9 | magenta | 498 | -0.471 | 1.5e-04 | *** | No | Yes | 1 | Has3 |
| 10 | purple | 421 | +0.132 | 0.315 | | No | No | 0 | - |
| 11 | greenyellow | 408 | -0.425 | 7.1e-04 | *** | No | Yes | 0 | - |
| 12 | tan | 403 | +0.228 | 0.080 | | No | No | 0 | - |
| 13 | salmon | 398 | -0.590 | 7.0e-07 | *** | No | Yes | 0 | - |
| 14 | cyan | 363 | +0.231 | 0.075 | | No | No | 0 | - |
| 15 | midnightblue | 360 | -0.648 | 2.1e-08 | *** | No | Yes | 1 | Hspg2 |
| 16 | lightcyan | 352 | -0.232 | 0.075 | | No | No | 0 | - |
| 17 | grey60 | 347 | -0.161 | 0.218 | | No | No | 0 | - |
| 18 | lightgreen | 318 | -0.151 | 0.251 | | No | No | 0 | - |

**Notes:** Modules ranked by gene count. Shading: Green = Top 6 by size; Blue = Trait-correlated only (p<0.05); Gold = Both criteria.

Module Selection Rationale: WGCNA modules were ranked by gene count (size), and the six largest modules per dataset were selected for detailed functional annotation to ensure adequate statistical power for pathway enrichment. Module-trait correlations were computed for all modules to identify injury-relevant transcriptional programs. Critically, the two selection criteria converge on the same biologically meaningful modules: among the Top 6 largest modules, those showing significant injury correlation ($p < 0.05$) contain the vast majority of predefined ECM axis genes. In BI, modules M2 (blue) and M4 (yellow) are both large and injury-correlated, collectively containing 21 ECM genes (81% of total). In SCI, modules M1 (turquoise), M2 (blue), and M5 (green) meet both criteria, with M1 and M5 containing 19 ECM genes (86% of total). This convergence validates the size-based selection approach while confirming that injury-correlated ECM programs are captured within the analyzed modules.

**Supplementary Figure S2. HA Turnover Gene Expression Analysis.**

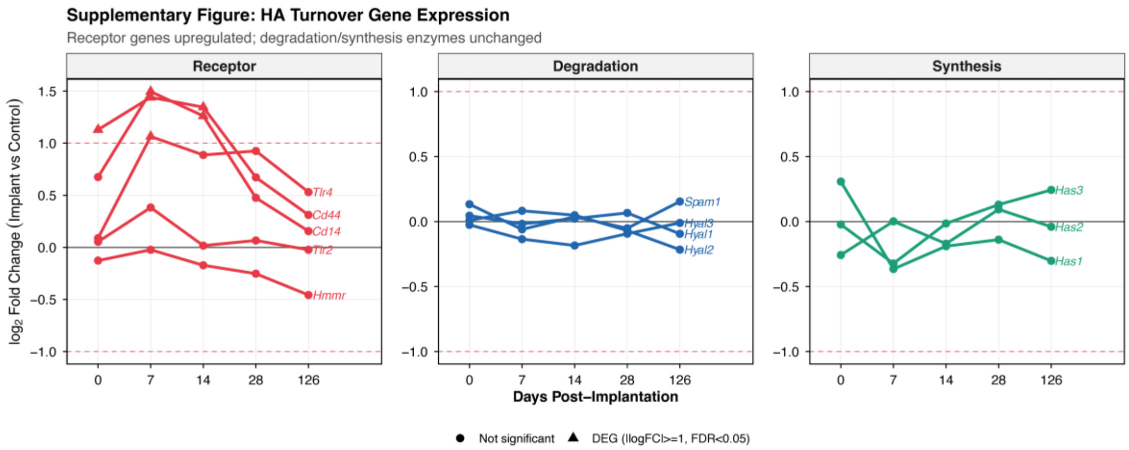


Line plots showing $\log_2$ fold change (Implant vs Control) for HA pathway genes across timepoints. **Left:** Receptor genes (*Cd44*, *Cd14*, *Tlr4*) showed significant upregulation peaking at Day 7, indicating enhanced HA fragment sensing. *Tlr2* and *Hmmr* showed no significant change, suggesting *Tlr4*/*Cd44* as the dominant sensing axis. **Center:** Degradation enzymes (*Hyal1-3*, *Spam1*) showed no significant changes at any time point. **Right:** Synthesis enzymes (*Has1-3*) showed no significant changes. The selective upregulation of receptors without corresponding changes in turnover enzymes suggests LMW-HA fragments were generated by initial insertion trauma rather than ongoing enzymatic degradation. Dashed red lines indicate DEG threshold ($\log_2 FC \geq 1$). Triangles indicate significant DEGs (FDR < 0.05).

#### Software environment

#### R pipeline (packages, versions, key parameters)

R 4.4.2. Core attached packages used in the pipeline (version):

Software Environment Discovery Analysis (BI/SCI Datasets - Mouse): R 4.4.2, WGCNA 1.73, limma 3.62.1, GEOquery 2.74.0, fgsea 1.32.2, clusterProfiler 4.14.4, org.Mm.eg.db 3.20.0, AnnotationDbi 1.68.0, mouse4302.db 3.13.0, pheatmap 1.0.12, ggplot2 3.5.1, tidyr 1.3.1, dplyr 1.1.4, plus Bioconductor infrastructure (Biobase 2.66.0, BiocGenerics 0.52.0, IRanges 2.40.1, S4Vectors 0.44.0) and helpers (fastcluster 1.2.6, dynamicTreeCut 1.63-1).

Validation Analysis (Neural Implant Dataset - Rat): R 4.3.0, oligo 1.62.2, limma 3.54.0, GSVA 1.46.0, fgsea 1.24.0, clariomsrattranscriptcluster.db (annotation package).

#### Brain Injury Modules

#### Module M1 (BI): Oxidative Metabolism and HA Degradation

Our results revealed a significant downregulation in Module M1 post-injury, aligning with secondary injury mechanisms known to contribute to neuronal death, with limited recovery observed by day 7. ROS production is particularly relevant to HA, as ROS can cleave the HA polymer (Stern et al. 2007; Duan und Kasper 2011; Yamazaki et al. 2003). ROS produced excessively after injury due to mitochondrial disruption can cleave HMW-HA into pro-inflammatory low molecular weight fragments (LMW-HA). These fragments exacerbate inflammation, activating innate immune responses and perpetuating secondary tissue damage.

Suppression of hub genes like GRIA1 and CAMK2A could underlie cognitive deficits observed after injury. Downregulation of myelin-related genes suggests compromised neural connectivity.
Additionally, key enzymes such as Hyal2, which regulate HA degradation, and collagen III (Col3a1) deposition likely influence scar stabilization while impeding axonal regeneration. Protecting neuronal integrity and maintaining ECM homeostasis may thus be essential strategies to mitigate the detrimental impacts identified in Module M1 after brain injury.

##### Module M2 (BI) : Innate Immune Activation and HA Fragment Signaling

A critical driver of the immune response observed in Module M2 is the role of low-molecular-weight hyaluronic acid (LMW-HA) fragments, generated after injury primarily through oxidative stress. These fragments act as damage-associated molecular patterns (DAMPs), binding TLR2 and TLR4 on microglia and macrophages and subsequently activating the MyD88-NF-κB pathway (Ruppert et al. 2014). This activation triggers the expression of pro-inflammatory cytokines such as IL-1β, TNF-α, IL-6, and IL-12, and chemokines, processes that are reflected in Module M2 gene signatures. This mechanism how LMW-HA engages TLRs leading to NF-κB activation and inflammatory gene expression.

parallels the immune activation seen with bacterial endotoxins like lipopolysaccharide (LPS), highlighting shared pathways such as NF-κB signaling.Thus, Module M2 likely includes genes encoding pattern-recognition receptors (TLRs), Cd14, NF-κB subunits, and inflammatory cytokines. The findings emphasize LMW-HA fragments' pivotal role in amplifying leukocyte recruitment and activation, contributing significantly to the secondary inflammatory wave following traumatic brain injury.

##### Module M3 (BI) : Cytokine Signaling and Extracellular Matrix Interaction

Although HA was not explicitly listed in the identified cytokine pathways, it critically modulates cytokine and growth factor signaling through interactions with its receptors, notably Cd44 (Fallacara et al. 2018). Following injury, Cd44 expression dramatically increases, amplifying its regulatory role over cytokine signaling essential for proliferation, migration, and gene expression. HA's interaction with Cd44 and other ECM components like neurocan and versican helps stabilize growth factor gradients, influencing cellular responses necessary for tissue repair. Particularly, HMW-HA acts as a crucial ECM element supporting regenerative, rather than inflammatory, processes.

Overall, Module M3 underscores how soluble cytokine signaling integrates with the HA-rich extracellular matrix via Cd44-mediated pathways to orchestrate cellular behaviors critical for post-injury healing and regeneration.

**Module M4 (BI) : Protein Synthesis and Innate Immune Regulation**

The integrated upregulation of protein synthesis and inflammatory pathways in Module 4 suggests that reactive astrocytes and microglia boost protein synthesis to support inflammation and extracellular matrix (ECM) production necessary during injury recovery. Critically, Module 4 likely regulates HA synthesis via HAS enzymes, essential for glial scar formation.

HA produced by reactive astrocytes aids tissue stabilization, modulates inflammation, and facilitates the transition from inflammatory (LMW-HA) to protective anti-inflammatory (HMW-HA) states. Moreover, active modulation of innate immunity through Tlr signaling by reactive glial cells characterizes this pivotal period. Ultimately, this dual role of ECM production and immune modulation helps stabilize tissue and manage inflammation in the subacute phase post-injury, significantly shaping recovery trajectories.

**Module M5 (BI): Vesicle Trafficking and Cell Communication**

Although Module 5's direct link to HA metabolism is less explicit than in other modules, it may still indirectly influence HA dynamics. Vesicle trafficking mechanisms contribute to HA metabolism by delivering and secreting precursor substrates and enzymes that regulate HA synthesis and degradation. Additionally, endocytic pathways mediated by vesicle transport are essential for internalizing and clearing HA fragments, which is critical for resolving inflammation.

Reactive astrocytes and microglia use secretory vesicles to release cytokines and matrix-modifying enzymes, indirectly affecting HA molecular weight dynamics and inflammatory states. Furthermore, synaptic vesicle cycling, potentially modulated by ECM-associated HA, may influence neuronal plasticity and excitability post-injury. HA degradation within synaptic perineuronal nets (PNNs) may initially enhance synaptic plasticity and neurotransmitter release, potentially exacerbating excitotoxicity, while subsequent HA restoration could stabilize synaptic connections.

Therefore, Module 5 represents critical cellular processes in vesicle-mediated trafficking that indirectly shape HA turnover and impact synaptic recovery dynamics following brain injury.

**Module M6 (BI): Stress Response and Metabolic Adjustment**

The biological processes captured in Module 6 point to cellular adaptations to injury-induced stress through metabolic reprogramming and oxidative stress management. HA metabolism intricately intersects these pathways: heightened ROS production after injury facilitates HA fragmentation into inflammatory forms (LMW-HA).

However, robust antioxidant responses emerging in the later phases could mitigate further HA breakdown, promoting a shift back toward stable HMW-HA-rich ECM environments conducive to healing. Additionally, chromatin remodeling pathways suggest epigenetic regulation may modulate gene expression relevant to HA metabolism, involving enzymes like HAS (synthesis) and hyaluronidases (degradation).

This dynamic regulation aligns with temporal shifts observed in HA-related gene expression and highlights the intricate interplay between metabolic stress management, HA turnover, and epigenetic adjustments during the later stages post-injury. Altogether, these processes contribute fundamentally to tissue stabilization, resolution of inflammation, and initiation of long-term repair following brain injury.

**SCI Modules**

**Module M1 (SCI): Synaptic Function and Neuroprotection**

Hyaluronic acid (HA) indirectly impacts Module M1 by influencing synaptic stability through perineuronal nets (PNNs), HA-rich extracellular structures that protect neurons. Following SCI, degradation of PNNs by oxidative stress and proteases can liberate synapses, increasing their vulnerability and potentially contributing to synaptic loss and aberrant plasticity.

Conversely, later re-establishment of HA-rich matrices may help support neuronal survival and functional recovery by stabilizing remaining synapses and modulating inflammation. Treatment with HMW-HA has demonstrated neuroprotective effects, limiting secondary damage and promoting synaptic recovery in SCI models.

Thus, Module M1 encapsulates the neuronal component of SCI, with HA playing a crucial, albeit indirect, role in governing synaptic resilience and the trajectory of functional recovery.

**Module M2 (SCI): Mitochondrial Activity and HA Turnover**

The ROS generated by SCI actively fragment hyaluronic acid (HA), shifting extracellular matrix (ECM) composition toward pro-inflammatory LMW-HA. This HA degradation exacerbates inflammation and secondary injury, reinforcing the cycle of oxidative damage.

In response, cellular adaptations include upregulation of antioxidant pathways and mitochondrial uncoupling mechanisms, aiming to minimize further oxidative stress and preserve matrix integrity. The continuous balance between HA degradation and synthesis is evident, with HA synthases potentially upregulated as a compensatory mechanism in response to oxidative damage.

Thus, Module M2 represents a metabolic adaptation to SCI that is intricately linked to dynamic changes in HA composition, reflecting the tissue's attempts to manage oxidative injury, limit inflammation, and promote eventual repair.

**Module M3 (SCI): Microglial Activation and Phagocytosis**

Hyaluronic acid (HA) plays a direct role in the clearance of fragmented extracellular matrix (ECM) debris; microglia and macrophages internalize HA fragments via Cd44 receptors, directing them to lysosomes for degradation. This HA-mediated phagocytosis promotes resolution of the initial inflammatory phase by removing pro-inflammatory debris. Additionally, HA fragments stimulate microglial activation through Tlr signaling, enhancing antigen-presenting capabilities and cytokine production, thereby potentially perpetuating inflammation.

Therefore, Module M3 encapsulates the phagocytic and immunoregulatory responses to SCI, critically modulated by HA dynamics. Effective clearance of HA fragments is pivotal, potentially determining whether inflammation resolves or persists chronically, ultimately impacting overall recovery potential.

**Module M4 (SCI): ECM Remodeling and Cell Signaling**

Relaxin signaling modulates fibrosis and ECM turnover, facilitating dynamic ECM remodeling crucial for mature scar formation. TNF signaling, significantly enriched in Module M4, influences astrocytes and fibroblasts, driving HA production and modulating Cd44 expression, further integrating inflammatory signals with ECM dynamics. HA plays a central role in Module M4's ECM remodeling processes.

After SCI, HA synthesis dramatically increases, creating a hydrated scaffold that supports other ECM components. Versican and aggrecan, major chondroitin sulfate proteoglycans (CSPGs), interact with HA to form aggregates stabilizing the glial scar but hindering axonal regeneration. HA indirectly interfaces with integrins via Cd44, mediating focal adhesion signaling and promoting astrocyte migration into injury sites. HA– Cd44 binding triggers PI3K-Akt signaling, enhancing cell survival and migration within scars.

Relaxin-induced ECM remodeling simultaneously promotes matrix hydration by increasing HA synthesis and collagen breakdown via matrix metalloproteinases (MMPs). Experimentally, HA synthesis inhibitors have been shown to reduce ECM deposition, suggesting HA's dual role: contributing to scar integrity while inhibiting full regenerative potential. Thus, Module M4 encapsulates the cellular programs underpinning chronic scar formation, orchestrated by HA's dynamic interactions with the ECM and surface receptors, ultimately stabilizing and isolating the lesion.

**Module M5 (SCI): Inflammation and Immune Response**

A key driver of Module M5 activation is low-molecular-weight hyaluronic acid (LMW-HA), produced by mechanical disruption and reactive oxygen species (ROS)-mediated fragmentation of high-molecular-weight HA (HMW-HA). LMW-HA fragments act as potent inflammatory mediators by engaging Toll-like receptors (TLRs), notably TLR2 and TLR4, on microglia, macrophages, and neutrophils. Activation of these receptors triggers downstream NF-κB signaling, amplifying inflammatory cytokine and chemokine production.
Cd44, a central HA receptor, plays dual roles—initially promoting inflammation by binding HA fragments, but later facilitating HA fragment clearance and inflammation resolution through interactions with regenerated HMW-HA.

Experimental interventions, such as exogenous delivery of HMW-HA or inhibitors of HA synthesis (e.g., 4-MU), have demonstrated the ability to modulate Module M5 activity, highlighting the importance of managing HA dynamics to prevent secondary damage and support tissue repair. Effective resolution of Module M5 activity depends on restoring HMW-HA dominance, thus shifting the tissue environment toward anti-inflammatory and regenerative states necessary for functional recovery.

**Module M6 (SCI): Delayed Repair, Lipid Metabolism, and HA Resolution**

This metabolic reprogramming aligned closely with the presence of lipid-rich "foam cell" macrophages observed chronically in SCI lesions, which ingest myelin-derived lipids and extracellular matrix debris, including glycosaminoglycans such as HA and CSPGs. Specifically regarding hyaluronic acid (HA), Module M6 significantly contributed to resolving HA-mediated inflammation.

By the chronic stage, early-generated LMW-HA fragments—known pro-inflammatory signals—were predominantly cleared, shifting towards a dominance of HMW-HA produced by reactive astrocytes.

Lysosomal pathways within macrophages facilitated the final breakdown of residual HA fragments into monosaccharides. Enzymes such as hexosaminidases and glucuronidases were central to this clearance process, consistent with the module's strong lysosomal enrichment. Importantly, Module M6 also signaled a transition to an anti-inflammatory phenotype, associated with M2-like macrophages, characterized by increased expression of regulatory genes like IL-10 and Arg1.

HMW-HA supported this transition by interacting with Cd44 receptors, promoting macrophage anti-inflammatory responses, and reducing pro-inflammatory cytokine production. Additionally, Module M6 indirectly fostered angiogenesis by removing inhibitory matrix elements and potentially producing angiogenic factors. Ultimately, SCI Module M6 encapsulated a critical reparative stage, resolving inflammation through lipid and glycan clearance, stabilizing the lesion into a chronic, metabolically stable, anti-inflammatory state marked by an HA-rich scar environment.

---

### Supplementary Code Appendix

All analysis scripts are publicly available at https://github.com/asharbatian/ECM_Axis_Neural_Injury_Analysis.

- Discovery analysis scripts (BI/SCI): discovery/ folder

- Validation analysis scripts: scripts/ folder

Note (applies to all scripts):

BI analyses omit LPS/Saline arms; figures display up to six modules. Parameters (e.g., power=6, bicor, signed TOM, minModuleSize=30, deepSplit=2, mergeCutHeight=0.2, reassignThreshold=0.8) and all file/folder names remain as in the analysis code.
